# LAMC2 marks a tumor-initiating cell population with an aggressive signature in pancreatic cancer

**DOI:** 10.1101/2022.03.24.485651

**Authors:** Donatella Delle Cave, Silvia Buonaiuto, Maria Mangini, Bruno Sainz, Annalisa Di Domenico, Tea Teresa Iavazzo, Gennaro Andolfi, Carme Cortina Duran, Marta Sevillano, Christopher Heeschen, Vincenza Colonna, Marco Corona, Antonio Cucciardi, Martina Di Guida, Eduard Batlle, Annachiara De Luca, Enza Lonardo

## Abstract

Tumor-initiating cells (TIC), also known as cancer stem cells, are considered a specific subpopulation of cells necessary for cancer initiation and metastasis. Here, we report a LAMC2-positive cell population, which is endowed with enhanced self-renewal capacity, and is sufficient for tumor initiation, differentiation, and driving metastasis. Mechanistically, mRNA profiling of these cells indicate a prominent squamous signature, and differentially activated pathways critical for tumor growth and metastasis, including deregulation of the TGF-β signaling pathway. Treatment with Vactosertib, a new small molecule inhibitor of transforming growth factor-β (TGF-β) type I receptor (activin receptor-like kinase-5, ALK5), completely abrogated the lung metastasis, primarily originating from LAMC2 expressing cells. Our results prompt further study of this TIC population in pancreatic cancer and exploration as a potential therapeutic target and/or biomarker.

## INTRODUCTION

Pancreatic ductal adenocarcinoma (PDAC) is the most common histologic subtype of pancreatic cancer. The disease ranks as the fourth leading cause of cancer-related deaths worldwide, with a dismal 5-year survival rate of less than 5% (Fitzmaurice and Global Burden of Disease Cancer Collaboration, 2018). PDAC is a devastating disease due to the late diagnosis, its aggressive nature (early metastasis), current lack of effective treatment options and a limited understanding of the cellular and molecular mechanisms underlying its initiation and progression. Furthermore, chemotherapy resistance and tumor recurrence are two unsettled problems associated with PDAC treatment (Gresham et al., 2014).

Many cancers including PDAC are hierarchically organized and constituted of functionally heterogeneous population of cells. Convincing evidence now demonstrates the critical relevance of the tumor-initiating cell (TIC) or cancer stem cell (CSC) compartment (Abel and Simeone, 2013). TICs shares several features with normal stem cells, such as self-renewal and differentiation potential (Lonardo et al., 2011; Cave et al., 2021). TICs bear crucial roles in tumor progression and metastasis (Hermann et al., 2007). From a clinical perspective, the principal issue with TICs relies on their refractory nature to conventional therapies, a feature that has been show to represent a main driver of tumor relapse (López-Gómez et al., 2016).

These data suggest that we need to reconsider our approach for diagnosing and treating cancer, as we should also take into consideration the successful elimination of TICs driving tumor progression. Thus, there is an urgent need to better understand the heterogeneous TICs subpopulations in order to develop more effective therapies against them. While several TIC markers have been proposed for PDAC (Li et al., 2007; Hermann et al., 2007; Bailey et al., 2014; Fox et al., 2016; Abel et al., 2018; Wang et al., 2019, 9; Cave et al., 2020), no one marker can accurately identify all TICs nor capture the diverse heterogeneity that exists within the TIC population. Consequently, there is a need to further study and characterize the TIC (sub-)populations in hopes of discovering new markers that can better define these populations. Here, we identify and characterize a TIC population in PDAC marked by high expression levels of the Laminin γ2 (LAMC2). LAMC2 is a subunit of the heterotrimeric glycoprotein laminin-332, which is a fundamental component of epithelial basement membranes and regulates cell motility and adhesion (Domogatskaya et al., 2012).

High LAMC2 expression correlates with poorer survival (Garg et al., 2014, 2). Using genetic CRISPR-Cas9 targeting, we show that LAMC2-high cells, compared with their CRISPR-edited LAMC2-low counterparts, display enhanced sphere-forming and *in vivo* tumorigenic capacities, enhanced migratory and invasive potentials and apparent chemoresistance.

RNA sequencing of LAMC2-high-generated tumors showed increased transforming growth factor beta (TGF-β) pathway activity and an aggressive squamous phenotype compared to the LAMC2-low-derived tumors. Specifically, we found that TGF-β/Smad signaling is essential for LAMC2-mediated maintenance of the TICs features and that LAMC2-low or KD (Knock-Down) reduced TGF-β/Smad signaling. Intriguingly, targeting the tumor bulk cells as well as LAMC2-high TICs with Vactosertib, a new specific inhibitor of the TGF-β receptor type 1 (TGFBR1), efficiently inhibited PDAC progression and abrogated metastasis. Therefore, this novel tumorigenic and metastatic LAMC2-high population in PDAC represents a promising new target for the development of more effective therapeutic strategies against this deadly disease.

## RESULTS

### LAMC2 correlates with poor outcome in PDAC patients

*LAMC2* transcriptional levels were evaluated using the publicly available transcriptome data sets Jandaghi (Jandaghi et al., 2016, 2), Janky (Janky et al., 2016), META (Martinelli et al., 2017, 6), and TCGA (Cerami et al., 2012). *LAMC2* mRNA levels were consistently and significantly higher in tumor tissue versus adjacent normal tissue (**Fig. 1a**). For the GSE21501 (Stratford et al., 2010), GSE28735 (Zhou et al., 2019), GSE62452 (Zhou et al., 2018) and GSE71729 (Moffitt et al., 2015) series, well-annotated clinical data are available and showed a significant decrease in median overall survival for *LAMC2* high-expressing patients compared to *LAMC2* low-expressing patients (**Fig. 1b**). Next, LAMC2 immunohistochemistry (IHC) was performed using a tissue microarray composed of 18 PDAC samples and three NP (Normal Pancreas) tissues (**Fig. 1c** and **Supplementary Fig. 1a**). LAMC2 expression was classified as 1 to 4 based on the *H*-score and stratified for PDAC stage 1 to 3, and the results revealed significantly increased LAMC2 levels in PDAC patients versus NP, and a clear correlation with disease progression (**Fig. 1d and Supplementary Fig. 1b**).

**Figure 1.**
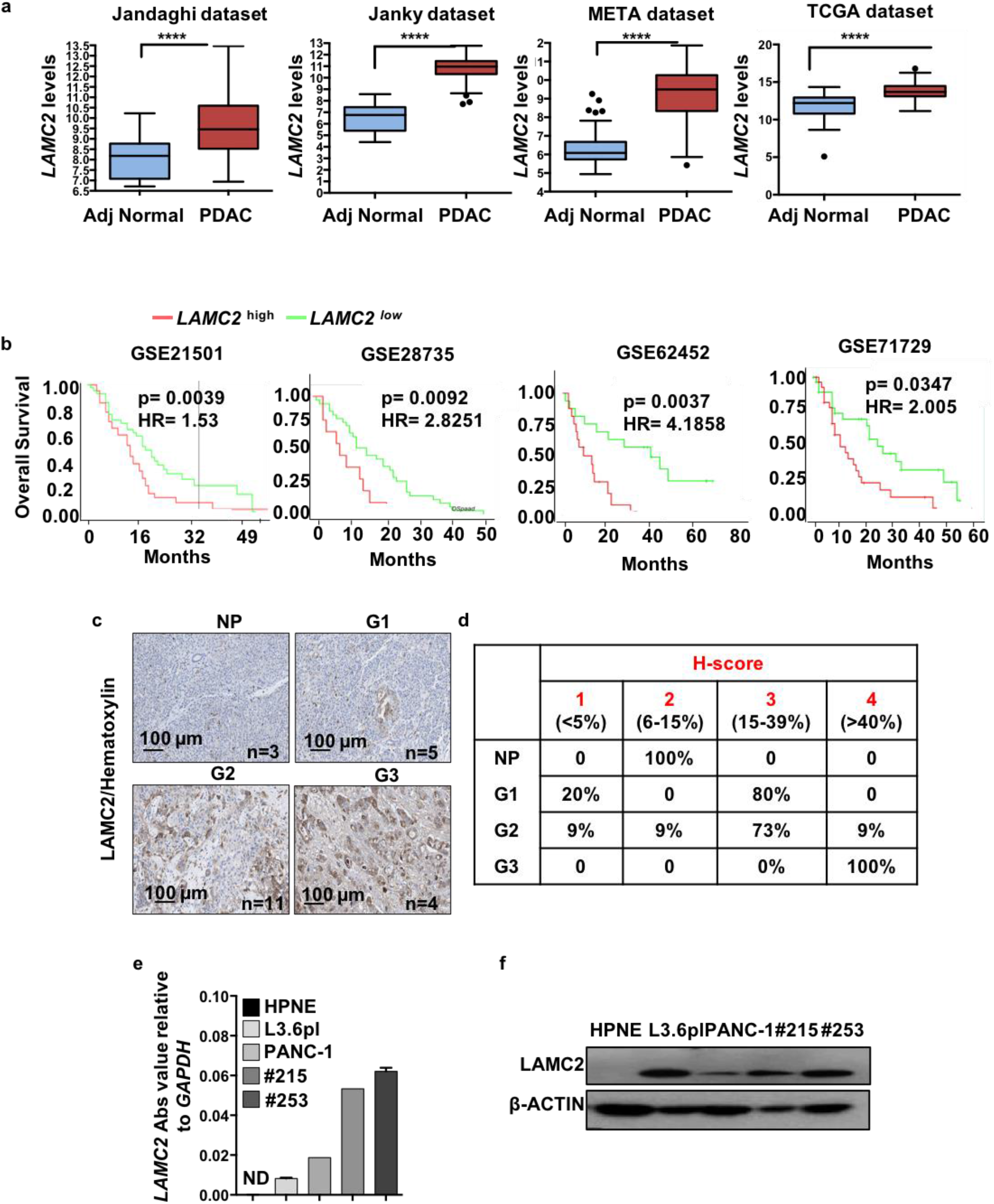
LAMC2 correlates with poorer outcome in PDAC patients. (**a**) Boxplots illustrating differential expression of *LAMC2* in PDAC tissue versus normal adjacent tissue using the indicated series of transcriptomic data. **** p < 0.0001. (**b**) Kaplan-Meier curves showing overall survival of PDAC patients, stratified according to the median value of *LAMC2* expression. (**c**) Representative images of IHC staining for LAMC2 (brown) in tissue sections from normal pancreas (P) and patients with PDAC tumors at G1, G2 and G3 grade. (**d**) *H*-score for LAMC2 expression. (**e**) qPCR analysis for *LAMC2* expression in adherent cells. Data are normalized to GAPDH expression. (**f**) Western blot analysis of LAMC2 in adherent cells. Parallel β-ACTIN immunoblotting was performed.

We also queried the Human Protein Atlas database for LAMC2 (Uhlen et al., 2010). A total of 12 patients with PDAC were classified based on LAMC2 for IHC staining, intensity and quantity. We observed that 9 out of 12 samples showed strong LAMC2 staining compared to 3 samples with medium levels of LAMC2; eleven samples presented strong (cytoplasmic/membranous) intensity for LAMC2 whereas one sample was classified as LAMC2 moderate; the distribution of stained cells showed one sample with <25%, two samples with 25– 75% and nine sample with >75% (**Supplementary Fig. 1c**). Notably, we also observed an association of LAMC2 expression with age and gender in the TMA (**Supplementary Fig. 1d-e**) and with age, alcohol and smoking in the TCGA (**Supplementary Fig. 1f)**. *In vitro*, we confirmed the absence of LAMC2 expression (gene and protein) in Normal Pancreatic cancer cells (i.e., HPNE (Campbell et al., 2007) compared to two established pancreatic cancer cell lines (L3.6pl and PANC-1) and two human PDAC-derived primary cultures (#215 and #253) (Lonardo et al., 2011; Rubio-Viqueira et al., 2006) (**Fig. 1e and Fig. 1f**). Together, these results suggest a clinical relevance of LAMC2 in PDAC.

### LAMC2 expression correlates with stemness

As poor outcome for PDAC has been related to the TIC/CSC content (Mizukami et al., 2014; Li et al.; Kim et al.), we hypothesized that higher expression of LAMC2 may be associated with stemness. First, we correlated the levels of LAMC2 (mRNA and protein) for cells cultured in adherent (Adh; enriched for differentiated cells) *versus* anchorage-independent conditions (Spheres, Sph; enriched for CSCs) (Lonardo et al., 2011). Quantitative real-time PCR (qPCR) (**Fig. 2a and Supplementary Fig. 2a**) and western blotting (**Fig. 2b and Supplementary Fig. 2b**) confirmed that LAMC2 was significantly upregulated in spheres compared to adherent cells. Concomitantly, stemness-associated genes (i.e., *CD44* and *CD133*) were overexpressed in spheres compared to adherent cells (**Supplementary Fig. 2c**). We recently demonstrated that the reduce expression of L1CAM, which we recently showed to represent a hallmark of stemness (Cave et al., 2020). Consistently, spheres also showed reduced *L1CAM* expression (**Supplementary Fig. 2c**). To exclude the possibility that the increased expression of *LAMC2* in spheres was related to differences in culture conditions (presence/absence of serum and plastic/no plastic), we compared adherent cells versus spheres by exchanging the media. We only observed *LAMC2* upregulation in cells grown as spheres (**Supplementary Fig. 2d**).

**Figure 2.**
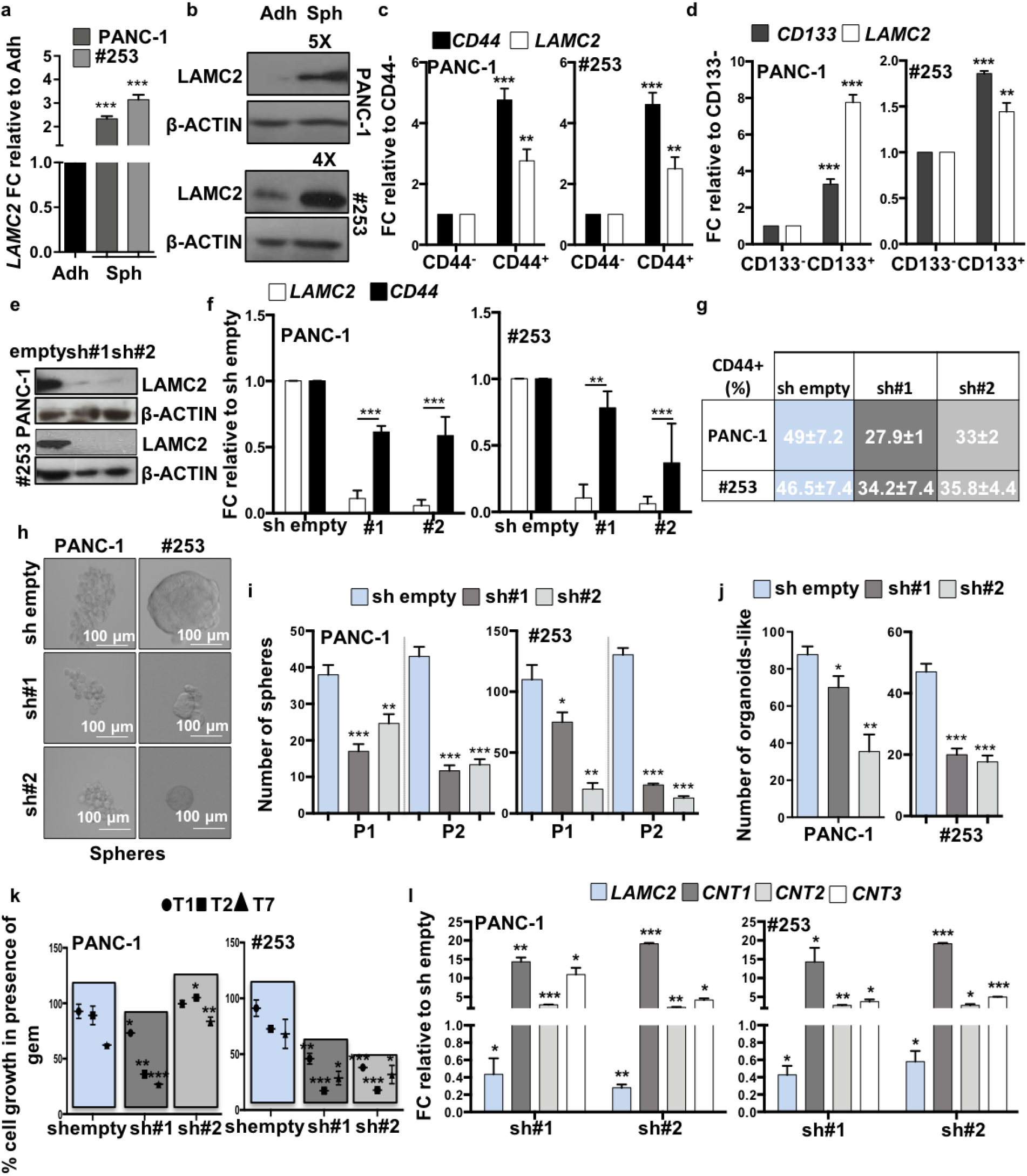
LAMC2 expression correlates with stemness and chemoresistance. (**a**) qPCR analysis for *LAMC2* gene expression in adherent cells versus spheres. Data are normalized to *GAPDH* and are presented as fold change in gene expression relative to adherent cells. (**b**) Western blot analysis for LAMC2 in adherent cells versus spheres. Parallel β-ACTIN immunoblotting was performed. (**c**) qPCR analysis for *CD44* and *LAMC2* gene expression in CD44^+^ sorted cells. Data are normalized to *GAPDH* and are presented as fold change in gene expression relative to CD44^-^ cells. (**d**) qPCR analysis for *CD133* and *LAMC2* gene expression in CD133^+^ sorted cells. Data are normalized to *GAPDH* and are presented as fold change in gene expression relative to CD133^-^ cells. (**e**) Western blot analysis for LAMC2 in sh empty and *LAMC2* knockdown cells. Parallel β-ACTIN immunoblotting was performed. (**f**) qPCR analysis for *LAMC2* and *CD44*expression for sh empty *versus LAMC2* knockdown cells. Data are normalized to *GAPDH* expression and are presented as fold change in gene expression relative to sh empty. (**g**) Flow cytometry quantification of CD44 in sh empty and *LAMC2* knockdown cells (**h**) Representative images of sh empty and *LAMC2* knockdown cells grown as spheres. (**i**) Sphere formation capacity of sh empty and *LAMC2* knockdown cells. P1= 1^st^ generation; P2= 2^nd^ generation. (**j**) Organoids formation capacity for sh empty *versus LAMC2* knockdown cells. (**k**) Growth capacity of sh empty and *LAMC2* knockdown cells in the presence of 100 μM of Gemcitabine (GEM). (**l**) qPCR analysis for *LAMC2, CNT1, CNT2* and *CNT3* genes in the sh empty *versus LAMC2* knockdown cells. Data are normalized to *GAPDH* expression and are presented as fold change in gene expression relative to sh empty. *p<0.05, **p<0.005, ***p<0.0005. n≥ 3.

Moreover, we tested the differentiation potential of the spheres as an important feature of cancer cell plasticity. For this purpose, we cultured L3.6pl, PANC-1 and #253 cells as spheres in the absence of serum for 7 days, and then plated them in adherent conditions in the presence of 10% FBS for 4 days. By qPCR we found that the expression of *LAMC2* was increased in spheres compared to the parental adherent cells and the levels were reduced in differentiated spheres (**Supplementary Fig. 2e**). At the same time, the expression of stemness genes (e.g., *ABCG2, CD133*, and *SOX2*) was significantly higher in spheres, compared to adherent cells, and the levels decreased in the differentiated spheres. Expectedly, *L1CAM* levels were lower in the spheres than in adherent, and its expression was restored in spheres upon differentiation (**Supplementary Fig. 2e**). Interestingly, the FACS-sorted CD44^high^ and CD133^high^ cells, respectively, showed a significant increase in *LAMC2* compared to their respective low/negative counterparts (**Fig. 2c-d and Supplementary Fig. 2f-g**). Lastly, *in vitro* treatment of L3.6pl, PANC-1, #215 and #253 with 100 μM of gemcitabine (Gem) revealed a significant increase in *LAMC2* levels in Gem-treated cells compared with vehicle treated cells (Ctrl) (**Supplementary Fig. 2h**). Taken together these data demonstrate that LAMC2 is enriched in the “standard” CSC population(s).

### Loss of LAMC2 affects stemness

To further corroborate above findings we next silenced *LAMC2* in L3.6pl, PANC-1, #215 and #253 using two different lentiviral shRNA constructs (sh#1 and sh#2) (**Fig. 2e-f and Supplementary Fig. 3a-b**). Upon knockdown of *LAMC2*, no apparent morphological changes were noted for cells cultured on plastic (**Supplementary Fig. 3c**).

While no changes in the cell cycle status were recorded, we observed modest alternations in cell viability (**Supplementary Fig. 3d-e**). Notably, the reduced expression of *LAMC2* did not translate into enhanced apoptosis (**Supplementary Fig. 4a**). By qPCR we observed that the sh*LAMC2* cells exhibited significant decreased mRNA levels for *CD44* compared with mock-infected cells (sh empty) (**Fig. 2f**), which could be confirmed by flow cytometry (**Fig. 2g and Supplementary Fig. 4b**). Modest changes were also observed in *CD133* mRNA (**Supplementary Fig. 4c**) or protein expression (**Supplementary Fig. 4d**). Consistently, we found that the number and size of both spheres (**Fig. 2h-i and Supplementary Fig. 4e-g**) and organoid-like structures (**Fig. 2j and Supplementary Fig. 4h**) were significantly decreased in sh*LAMC2* cells compared to sh empty cells.

Next, we tested the knockdown cells for chemoresistance and found that sh*LAMC2* cells were more sensitive to gemcitabine treatment than the sh empty cells (**Fig. 2k**). Gemcitabine is transported by multiple active nucleoside transporters (e.g. *CNT1, CNT2* and *CNT3*) and we found augmented expression for all of them in sh*LAMC2* cells compared to sh empty cells (**Fig. 2i**). Transmigration (**Fig. 3a-b and Supplementary Fig. 4i-j**), wound healing (**Supplementary Fig. 4k**) and gelatin degradation (**Fig. 3c-d and Supplementary Fig. 5a-b**) assays confirmed that sh*LAMC2* cells were less aggressive compared to sh empty cells. Of note, as shown above (**Supplementary Fig. 3d**), we did not observe decreased viability in sh*LAMC2* cells compared with sh empty cell, thereby excluding that the smaller number of transmigrated sh*LAMC2* cells was merely due to their reduced proliferation. Tumor cell invasion requires loss of cell–cell interactions, and is often associated with the epithelial–to-mesenchymal transition (EMT). qPCR analysis revealed upregulation of epithelial marker *CDH1* (E-Cadherin), whereas the mesenchymal transcription factors (i.e. *SNAIL1* and *VIMENTIN*) were downregulated in sh*LAMC2* cells compared with sh empty cells (**Fig. 3e and Supplementary Fig. 5c-d**). Increased matrix metalloproteases (MMPs) levels are, nowadays, considered as important sign of the pro-tumorigenic and invasive potential of tumor cells. In line with the aforementioned observations, we observed a significant reduced expression of *MMP2* and *MMP10* in sh*LAMC2* cells compared with sh empty cells (**Fig. 3f and Supplementary Fig. 5e**). Lastly, we subcutaneously injected 250,000 L3.6pl, PANC-1, #215 or #253 sh empty or sh*LAMC2* cells into nude mice and observed that tumors derived from sh*LAMC2* cells formed later, were less and were smaller compared with sh empty cells (**Fig. 3g and Supplementary Fig. 5f**). Importantly, qPCR analysis confirmed that *LAMC2* was still downregulated in the sh*LAMC2* tumors, and that they exhibited reduced levels of mesenchymal and metalloprotease (**Supplementary Fig. 5g**) genes as well as non-detectable (for PANC-1) or significantly reduced levels of *CD44* cell surface expression (**Fig. 3h**).

**Figure 3.**
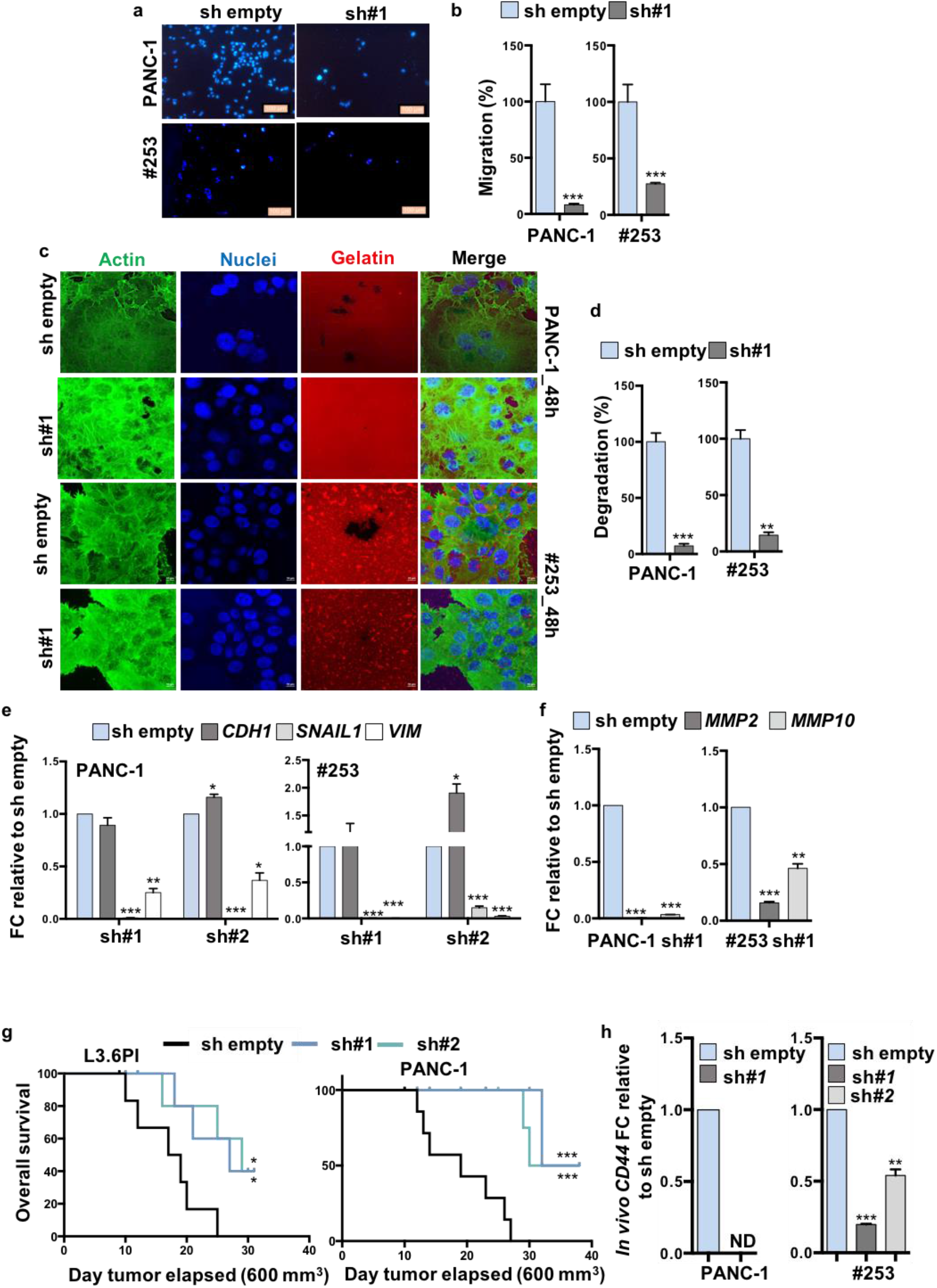
Loss of LAMC2 diminishes tumorigenicity. **(a)** Migration assay for sh empty *versus LAMC2* knockdown cells. The nuclei were stained with DAPI (blue). **(b)** Migratory potential of sh empty *versus LAMC2* knockdown cells. (**c**) Representative images of gelatin degradation for sh empty *versus LAMC2* knockdown cells. Nuclei were stained with Hoechst 33342 (blue), green represents actin (Alexa Fluor(tm) 488 Phalloidin) and red illustrates gelatin (Rodhamine). (**d**) Invasive potential of sh empty *versus LAMC2* knockdown cells. (**e**) qPCR analysis for EMT genes in sh empty and *LAMC2* knockdown cells. Data are normalized to *GAPDH* and are presented as fold change in gene expression relative to sh empty. (**f**) qPCR analysis for *MMP2* and *MMP10* gene expression in sh empty and *LAMC2* knockdown cells. Data are normalized to *GAPDH* and are presented as fold change in gene expression relative to sh empty. (**g**) Kaplan–Meier curve for sh empty and *LAMC2* knockdown cells subcutaneously xenografted into athymic CD1 mice. n≥ 10. (**h**) qPCR analysis for *CD44* gene expression in sh empty and *LAMC2* knockdown cells isolated from respective tumors. Data are normalized to *GAPDH* and are presented as fold change in gene expression relative to sh empty. *p<0.05, **p<0.005, ***p<0.0005. n≥ 3.

### Generation of *LAMC2*-EGFP knock-in human pancreatic cancer cells

LAMC2 is a secreted molecule present in the ECM of the cells, which rendered sorting of the cells based on LAMC2 expression and their subsequent downstream analysis difficult. We thus designed a strategy using CRISPR/Cas9-mediated homologous recombination to mark *LAMC2* cells. For these experiments, we selected L3.6pl, PANC-1 and #253 cells. The targeting strategy is summarized in **Figure 4a** and detailed in the Materials and Methods section. In brief, we designed Cas9 single-guide RNA complementary to sequences overlapping the stop codon of the *LAMC2* locus and generated a donor vector that contained *LAMC2* homology arms flanking an EGFP reporter cassette positioned immediately upstream of the stop codon. We added an LF2A self-cleavage peptide (de Felipe et al., 2010) in frame with EGFP so that the *LAMC2*-EGFP locus would be transcribed as a single mRNA, whereas the resulting polypeptide would be cleaved in the two encoded proteins, LAMC2 and EGFP (**Fig. 4a**). Next, we nucleofected the L3.6pl, PANC-1 and #253 cells with the donor vector together with a guide-RNA-Cas9 (guide), and after 48 hours we sorted cells that had incorporated the sgRNA-Cas9 px330 vector, which had an IRFP selectable cassette incorporated (Cortina et al., 2017) (IRFP^+^ cells) (**Supplementary Fig. 6a**). About 26-31% in L3.6pl, 33% in PANC-1 and 4-6% in #253 IRFP^+^ cells expressed EGFP after 20 days in culture (**Supplementary Fig. 6b**). Subsequently, we generated single cell-derived cultures and assessed the specific integration of the EGFP reporter cassette by PCR (**Supplementary Fig. 6c**). These analyses showed that 100% of L3.6pl, 100% of PANC-1 and 37.5% (3 out of 8) of #253 clones had correctly integrated the EGFP reporter in the *LAMC2* locus. In these single cell-derived knock-in cell cultures, every culture was composed of an admixture of cells expressing distinct EGFP levels (**Supplementary Fig. 6d**). *LAMC2*-EGFP^+^ cells isolated by FACS expressed highest *LAMC2* mRNA levels confirming that EGFP correctly reported the endogenous LAMC2 expression (**Fig. 4b and Supplementary Fig. 6e**).

**Figure 4.**
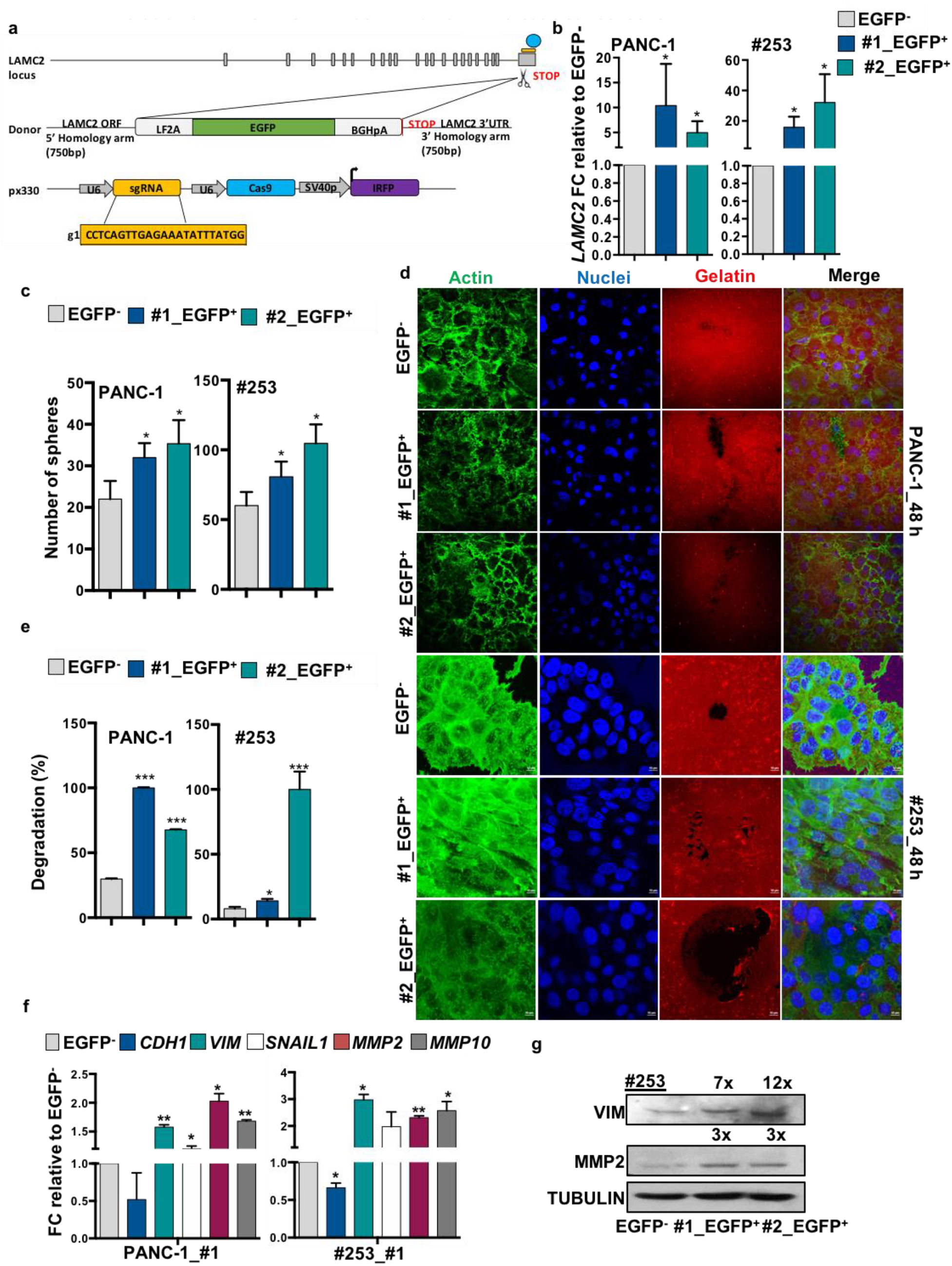
Generation of *LAMC2*-EGFP knock-in human pancreatic cancer cells. (**a**) Design of LAMC2-EGFP donor and CRISPR/Cas9 sgRNA vectors. Blue circle represents the CRISPR/Cas9 protein complex and the yellow box underneath illustrates the guide RNA. (**b**) qPCR analysis for *LAMC2* gene expression in EGFP^+^ and EGFP^-^ cells. Data are normalized to *GAPDH* and are presented as fold change in gene expression relative to the EGFP^-^ counterpart. (**c**) Sphere formation capacity for EGFP^+^ *versus* EGFP^-^ cells. (**d**) Representative images of gelatin degradation for EGFP^+^ *versus* EGFP^-^ cells. Nuclei were stained with Hoechst 33342 (blue), green represents actin (Alexa Fluor(tm) 488 Phalloidin) and red illustrates gelatin (Rodhamine). (**e**) Invasive potential of sh empty *versus LAMC2* knockdown cells. (**f**) qPCR analysis for EMT and *MMP2* and *MMP10* gene expression for EGFP^+^ *versus* EGFP^-^ cells. Data are normalized to *GAPDH* and are presented as fold change in gene expression relative to the EGFP^-^ counterpart. (**g**) Western blot analysis of VIM and MMP2 in EGFP^+^ and EGFP^-^ cells. Parallel Tubulin immunoblotting was performed. *p<0.05, **p<0.005, ***p<0.0005. n≥ 3.

### Characterization of human *LAMC2*-EGFP PDAC cells *in vitro* and *in vivo*

We next investigated the properties of the LAMC2-EGFP^+^ cells. Morphologically we did not observe, as expected, significant changes in the LAMC2-EGFP-clones (EGFP^+^#1 and EGFP^+^#2) compared to control cells (EGFP^-^) (**Supplementary Fig. 6f**). Likewise, the cell cycle status was essentially unaltered among LAMC2-EGFP-clones and EGFP^-^ cells (**Supplementary Fig. 6g**). To confirm that the LAMC2-EGFP^+^-clones maintained their intrinsic CSC properties, we examined their ability to grow as spheroids in standard sphere-forming medium. After 7 days of culture, the numbers of spheres formed were greater among LAMC2-EGFP^+^-clones versus the corresponding EGFP^-^ cell populations (**Fig. 4c and Supplementary Fig. 6h**). Moreover, the size (but not the number) of organoid-like structures was significantly increased in LAMC2-EGFP^+^-clones compared with EGFP^-^ cells (**Supplementary Fig. 7a-c**). Gelatin degradation assay also confirmed that LAMC2-EGFP^+^-clones were more aggressive compared with EGFP^-^ cells (**Fig. 4d-e**). qPCR analysis showed that LAMC2-EGFP^+^-cells indeed exhibited increased expression levels for mesenchymal genes (*VIM* and *SNAIL1*) and metalloproteases genes (*MMP2* and *MMP10*), respectively, compared to levels for the corresponding EGFP^-^ cells (**Fig. 4f-g and Supplementary Fig. 7d**).

Next, we subcutaneously inoculated CD1 nude mice with LAMC2-EGFP^+^ or EGFP^-^ cells and investigated their *in vivo* tumorigenicity. LAMC2-EGFP^+^-cells were able to generate earlier and bigger tumors compared to EGFP^-^ cells (**Fig. 5a and Supplementary Fig. 7e**). Histological analysis revealed that the tumor xenografts derived from LAMC2-EGFP^+^-clones displayed a glandular organization and increased desmoplasia compared with tumor xenografts derived from EGFP^-^ cells (**Fig. 5b and Supplementary Fig. 7f**). A substantial proportion of the epithelial compartment of the tumor showed LAMC2-EGFP^+^ cells, with their EGFP levels varying considerably (**Supplementary Fig. 7f-g**). Tumors generated from LAMC2-EGFP^+^ cells were populated with LAMC2-EGFP^+^ and EGFP^-^ tumor cells thus implying that LAMC2-expressing cells undergo self-renewal and differentiation during tumor formation and expansion (**Supplementary Fig. 7f-g**). Of note, xenografts generated by EGFP^-^ cells contained both LAMC2-EGFP^+^ and EGFP^-^ cells (**Supplementary Fig. 7f**), indicating plasticity. By qPCR analysis we determined that tumors derived from LAMC2-EGFP^+^ cells expressed more than 5-fold higher levels of *LAMC2* than tumors derived from EGFP^-^ cells (**Fig. 5c and Supplementary Fig. 7h**). Moreover, LAMC2-EGFP^+^ cells were enriched for *VIM, SNAIL1, MMP2* and *MMP10* genes (**Fig. 5c and Supplementary Fig. 7h-i**). Interestingly, EGFP^+^ cells displayed enhanced expression of CSC markers *CD44* and *CD133* compared to EGFP^-^ cells (**Fig. 5d**). The expression levels for the aforementioned genes were comparable to levels observed *in vitro*. To assess the capacity of these tumor cell populations to serially propagate the disease into secondary hosts, we inoculated 200 or 1,000 LAMC2-EGFP^+^ or EGFP^-^ epithelial tumor cells into new mice. These experiments showed that the LAMC2-EGFP^+^ cell population was strongly enriched for TIC cells compared to their differentiated EGFP^-^ counterparts (**Fig. 5e**). Specifically, the LAMC2-EGFP^+^ fraction from L3.6pl cells formed tumors with 200 cells (2/8) while the EGFP^-^ fraction formed only very small tumors with 1,000 cells (4/8). For the PANC-1 cell line, 200 isolated LAMC2-EGFP^+^ cells initiated tumors (6/8) whereas 1,000 cells were required to initiated tumors in the EGFP^-^ (4/8) group (**Fig. 5e**).

**Figure 5.**
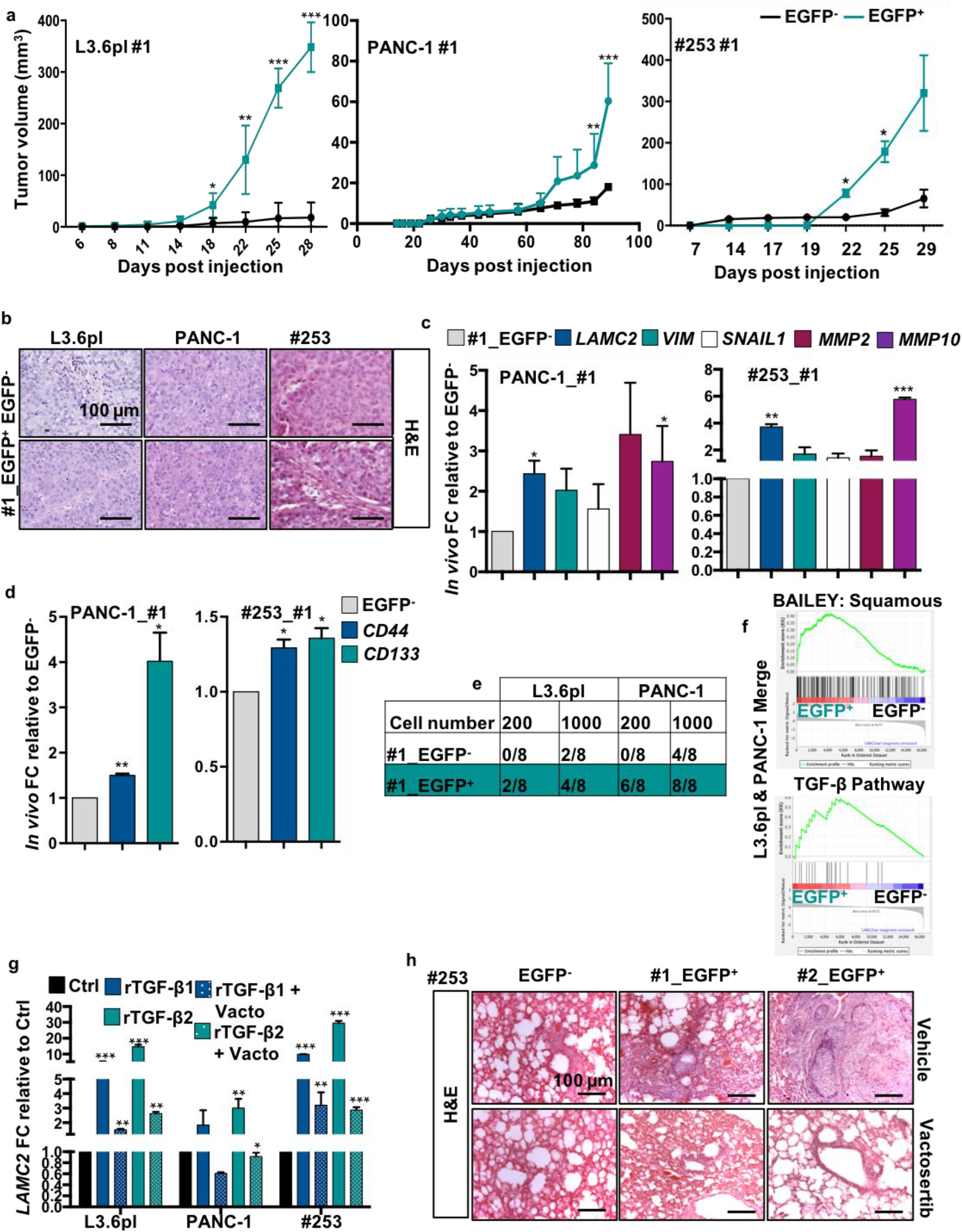
Pharmacological inhibition of TGF-β signaling blocks LAMC2-induced metastasis. (**a**) Tumor volume of EGFP^+^ and EGFP^-^ cells subcutaneously injected into athymic CD1 mice. n ≥ 10. (**b**) Representative H&E-stained sections of xenografts derived from EGFP^+^ and EGFP^-^ cells, respectively. (**c**) qPCR analysis for *LAMC2*, EMT, *MMP2* and *MMP10* gene expression in EGFP^+^ and EGFP^-^ cells, respectively, isolated from tumors. Data are normalized to *GAPDH* and are presented as fold change in gene expression relative to EGFP^-^ (**d**) qPCR analysis for *CD44* and *CD133* expression in EGFP^+^ and EGFP^-^ cells, respectively, isolated from tumors. Data are normalized to *GAPDH* and are presented as fold change in gene expression relative to EGFP^-^. (**e**) Number of tumors generated by subcutaneous injection of EGFP^+^ and EGFP^-^ cells, respectively. (**f**) Enrichment plot for EGFP^+^ *versus* EGFP^-^ cells isolated by FACS from subcutaneous tumors. (**g**) qPCR analysis for *LAMC2* expression in PDAC cells treated with 10 ng/ml of rTGF-β1 in the presence or absence of 80 μM Vactosertib. Data are normalized to *GAPDH* and are presented as fold change in gene expression relative to control. (**h**) Representative H&E-stained sections of lungs following tail vein injection of EGFP^+^ and EGFP^-^ tumor cells. Mice were treated with Vactosertib (40 mg/kg mice) or vehicle. *p<0.05, **p<0.005, ***p<0.0005. n≥ 3.

### Transforming growth factor beta (TGF-β) signaling inhibitor blocks LAMC2-induced metastasis

To identify key cellular pathways defining the specific features of LAMC2 expressing cells, transcriptomic analysis was performed using RNA-seq on LAMC2-EGFP^+^ and EGFP^-^-derived tumors (**Fig. 5f and Supplementary Fig. 8**). Gene set enrichment analysis (GSEA) revealed that pro-metastatic signatures represent a hallmark of LAMC2-EGFP^+^ -derived tumors. Interestingly, we observed that the LAMC2-EGFP^+^ -derived tumor shown significant enrichment in the squamous subtype transcriptional profile (**Fig. 5f**), which has been linked to hypermethylation and down-regulation of genes determining endodermal identity in the pancreas, and is associated with poor outcome (Australian Pancreatic Cancer Genome Initiative et al., 2016). Moreover, we found enrichment for transcripts related to the TGF-β (**Fig. 5f**), focal adhesion, hypoxia, and MAP Kinases pathways (**Supplementary Fig. 8 and 9a**). Encouraged by above results we next examined a potential link between LAMC2 expression and TGF-β/Smad signaling. Notably, *in vitro* treatment of PDAC cells with recombinant TGF-β1 or TGF-β2 strongly increased LAMC2 expression (**Fig. 5g and Supplementary Fig. 9b**), which was significantly reduced by the co-treatment with Vactosertib, a new clinical-grade inhibitor of TGF-β receptor I (**Fig. 5g**). Metastatic PDAC is caused by disseminated cancer cells that maintain the capacity to initiate new lesions in distant tissues (Yachida et al., 2010), which for PDAC are mainly liver and lung. To assess the ability of the LAMC2-EGFP^+^ cells to generate metastases, we inoculated them as dissociated cells via the tail vein of nude CD1 mice. Even thousands of injected EGFP^-^ tumor cells produced hardly any metastases (**Fig. 5h**), whereas EGFP^+^ cells were highly metastatic (**Fig. 5h**). Intriguingly, treatment of mice with Vactosertib completely abrogated the capacity of LAMC2-EGFP^+^ cells to form metastasis (**Fig. 5h and Supplementary Fig. 9c**), underlying that the TGF-β/Smad signaling pathway is a key driver of LAMC2^+^ CSC-mediated metastases in vivo.

## DISCUSSION

TIC or CSC have been proposed to represent key drivers for tumor progression and metastasis formation in PDAC. However, as of today, there are still no effective and translatable therapeutic strategies that actually eradicate this subset of highly aggressive cancer cells. Efficiently targeting TICs appear to be essential to achieve cure, due to their self-renewal capacity and resistance to conventional chemotherapies (Zhou et al., 2021), and therefore they are the drives tumor relapse. Despite significant advances in our understanding of TIC biology, the identification of specific markers to help isolate these cells remains largely debated, incompletely established, and limited to a handful of markers that are neither solely expressed on TICs nor expressed across all TIC subpopulations. Here, we demonstrated that PDAC cells with high expression of LAMC2 were highly metastatic, consistent with the inverse correlation between LAMC2 expression and patient survival.

LAMC2 levels have been already associated with poor prognosis in several human cancers due to its effects on cancer cell proliferation, migration and invasion. In pancreatic cancer, by performing both integrated bioinformatics analysis and proteomic assays, its potential as a new putative biomarker has emerged (Islam et al., 2021). Of note, elevated serum levels of LAMC2 in patients with pancreatic cancer make it an attractive serum-based diagnostic marker (Kosanam et al., 2013). Nevertheless, no evidences regarding LAMC2 in driving the stem properties and the metastatic capacities of PDAC cells have been reported.

Here, we showed that the knockdown of *LAMC2* resulted in decreased self-renewal, migration, invasion, tumorigenicity and chemoresistance and it is reflected *in vivo* by a reduced tumorigenic potential, thus suggesting that silencing of LAMC2 counteracts the TIC phenotype and PDAC aggressiveness. The CRISPR/Cas9 technology allowed us to study of human tumors through genetic manipulations that had only been feasible to date in animal models. This technological advance is particularly well suited to analyze phenotypic diversity of cell populations within cancers as it enables genetic editing of genes. For example, the labelling of distinct tumor cells with specific marker genes, which are not necessarily expressed at the cell surface such as LAMC2. Therefore, tumors generated from edited cells reflect the behaviour of a single TIC lineage in a genetically homogenous mutational background. For the first time, this approach now enabled us to explore the behavior of LAMC2-EGFP^+^ cells in intact tumors, and helped us to demonstrate directly how this cell population contributes to growth and dissemination in PDAC.

First, we showed that LAMC2-EGFP^+^ cells displayed EMT molecular traits concomitant with an increased migratory activity, which explained the enhanced metastatic features of these cells when injected in immunocompromised mice injected. Transcriptomic analysis revealed a signature of LAMC2-EGFP^+^ cell-derived tumors consistent with the squamous subtype as defined in the Bailey classification (Australian Pancreatic Cancer Genome Initiative et al., 2016). In this classification, squamous is the molecular subtype of PDAC with the worse prognosis. In addition, enrichment of gene networks involved in inflammation, hypoxia, metabolic reprogramming, TGF-β signaling, MYC pathway activation and its target genes were also observed and are key features of the squamous PDAC tumors.

Interestingly, the activation of TGF-β signaling plays a key role in promoting TIC migration and metastasis (Lonardo et al., 2011; Cave et al., 2020). Cytokines released in the tumor microenvironment significantly contribute to maintain the undifferentiated state and clonogenic activity of TIC (Truong and Pauklin, 2021). Specifically, here we found TGF-β1 to be a potent inducer of LAMC2 expression. *In vivo*, a similar effect might be induced by TGF-β1 derived from stromal and/or LAMC2-high cells.

While stromal cells seem to play a key role in the production of TGF-β1 (Grauel et al., 2020), it is likely that autocrine production of TGF-β1 may contribute to promoting the metastatic activity of LAMC2-EGFP^+^ cells. Intriguingly, Vactosertib completely prevented metastasis formation by LAMC2-EGFP^+^ cells. Vactosertib is a novel and orally administered transforming growth factor-β (TGF-β) type I receptor inhibitor that is currently in clinical testing for other cancer types (https://www.clinicaltrials.gov/ct2/show/NCT04258072). Therefore, our findings may have important therapeutic implications for PDAC. Although some LAMC2-low cells also bear modest tumorigenicity and are capable of giving rise to small tumors, the vast majority of tumorigenic capacity seems to be confined to the LAMC2-high population, which contains virtually all the cells with high level of TGF-β activation. Because tumorigenic capacity is a prerequisite to successfully seed metastatic lesions, it is likely that the relative density of TICs in PDAC directly correlates with the metastatic propensity of tumor lesions. In this context, LAMC2 could serve as a key functional biomarker that is critically involved in promoting the main features of cancer progression.

## MATERIALS AND METHODS

**PDAC cultures**. Tumor-derived primary cell lines #215 and #253 (**Tissue derivation**: primary pancreatic tumor; **Carcinoma type**: pancreatic ductal adenocarcinoma) were cultured in RPMI, 10% FBS, and 50 units/ml penicillin/streptomycin (Lonardo et al., 2011). The human PDAC cancer cell lines L3.6pl (**Tissue derivation**: metastatic lymph node; **Carcinoma type**: adenosquamous carcinoma), PANC-1 (**Tissue derivation**: pancreatic tumor; **Carcinoma type**: ductal carcinoma) and immortalized HPNE were maintained as previously described (Lonardo et al., 2011). Their identity (annually) and *Mycoplasma* free-state (bi-weekly) is routinely tested by DNA fingerprinting using short tandem repeat (STR) profiling, and by PCR-based, MycoAlert Mycoplasma Detection Kit (Lonza, Bioscience). Each cell lines were used within passage 4/5 since their thawing from originally frozen vials.

### RNA Preparation and Real-Time quantitative PCR

Total RNAs were extracted with Eurogold TRIFAST kit (Euroclone) according to the manufacturer’s instructions. One microgram of total RNA was used for cDNA synthesis with High-Capacity reverse transcriptase (Thermofisher). Quantitative real-time PCR was performed using SYBR Green PCR master mix (Thermofisher), according to the manufacturer’s instructions. The list of utilized primers is depicted in **Table 1**.

**Table 1.**
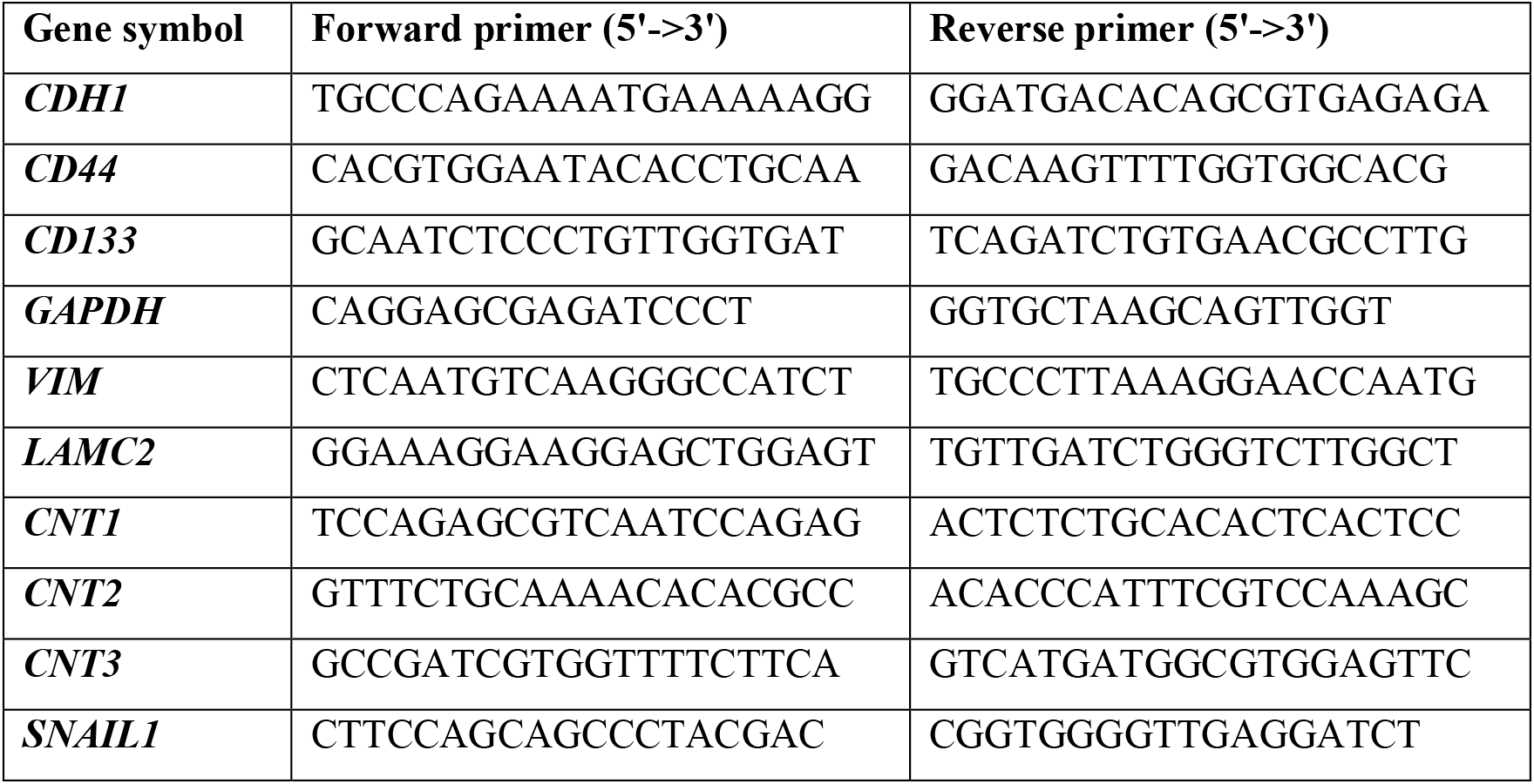
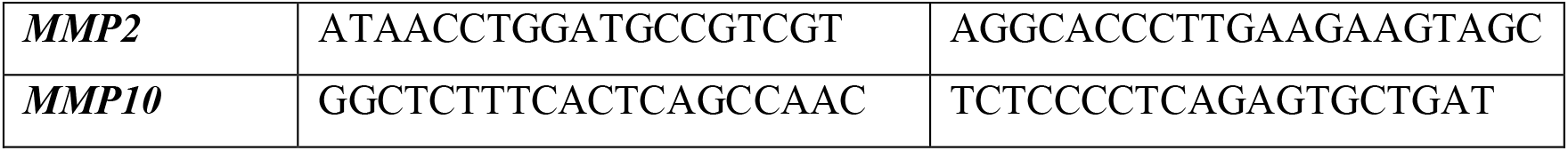

### Protein isolation and western blot analysis

Cells were lysed with RIPA buffer (50mM Tris-HCl at pH 7.6, 150mM NaCl, 1% NP-40, 0.5% sodium deoxycholate, 0.1% SDS, 5 mM EDTA plus proteases and phosphatases inhibitors) for 1 hour at 4^°^C. Total protein quantification was performed with Bio-Rad Protein Assay Dye Reagent concentrate. A total of 40 μg of protein was separated on 15% SDS–PAGE gels at 100 V and transferred to PVDF membranes for 2 hours at 200 mA. PVDF membranes were hybridized with antibodies against LAMC2 (AMAB 91098, Sigma-Aldrich), β-actin (E-AB-20058, Elabscience), MMP-2 (4022S, Cell Signaling), Vimentin (3932, Cell Signaling) and α-tubulin (2144, Cell Signaling) treated with peroxidase-conjugated goat anti-mouse or anti-rabbit Ig secondary antibody (DPVR-HRP, Immunologic), and then visualized by enhanced chemiluminescence (ECL Nova 2.0 XLS071, 2050 Cyanagen). n>6.

### Lentiviral shRNA delivery

As lentiviral shuttle backbone we used a pLKO shRNA plasmid (Mission SIGMA). As control we used pLKO shRNA empty expression vectors. Cells were then transduced with lentiviral particles in the presence of polybrene (8 µg/ml, Sigma). The cells were seeded at a density of 30,000 cells per well in a 24 multiwell plates and allowed to adhere overnight. The next day, the cells were infected with the lentiviral particles for 6 hours. Stably transduced cells were obtained using puromycin resistance.

### Flow cytometry and cell sorting

Flow cytometry analysis or flow cytometry cell sorting was performed using anti-human CD133-APC (BioLegend – 372805) and anti-human CD44-APC (BioLegend – 338805) to identify CSCs. 7AAD (BD) was used for exclusion of dead cells. Samples were analyzed by flow cytometry using a FACS Canto II (BD) and data were analyzed with FlowJo 9.2 software (Ashland, OR). Experiments were repeated a minimum of three times independently, in triplicate samples.

### Immunofluorescence

Cells were fixed in 4% paraformaldehyde (PFA) for 20 min at room temperature. After blocking with 5% bovine serum albumin in PBS-Triton 0.1%, cells were incubated with unconjugated primary antibody: LAMC2 (SIGMA, HPA024638-100U) overnight at 4^°^C in the dark, and the day after counterstained with fluorescent secondary antibody. The nuclei of cell were stained by incubating with DAPI (SIGMA). Images were acquired at room temperature using the LEICA DMI6000 inverted microscope (Leica, Heidelberg, Germany) on a DC 350 FX camera (Leica).

### Sphere-formation assay

Pancreatic cancer spheres were generated and expanded in CSCs media composed by: Advanced DMEM:F12 (GIBCO) supplemented with 1× glutaMAX (GIBCO), 1× B-27 (GIBCO), 1× N2 (GIBCO), 20 ng/ml bFGF (basic fibroblast growth factor) (Invitrogen), and 50 ng/ml EGF (epidermal growth factor) (Peprotech, London, UK). Five hundred cells per 500µl of sphere medium were seeded in 24 ultra-low attachment plates (Corning B.V., Schiphol-Rijk, Netherlands) as described previously (Lonardo et al., 2015). After 7 days of incubation, spheres were typically > 75 µm large. For serial passaging, 7-day-old spheres were harvested using 40 µm cell strainers, dissociated to single cells with trypsin, and then regrown for 7 days. Cultures were kept no longer than 4 weeks after recovery from frozen stocks (passage 3-4).

### Matrigel embedding culture assay

Five hundred of PDAC cells were embedded in 50 µL of 100% BME2 (Cultrex) and seeded in 24 multiwell plates (Corning). The formed spheres, here termed organoid-like structures, were cultured in CSCs media for 7 days.

### Migration assays

Migration assays were performed using Boyden chambers (Corning). 25,000 tumor-derived primary cell lines #215 and #253 and PANC-1 (sh empty or sh*LAMC2)* were added to the inserts of the chamber for 22 hours at 37^°^C. Migrated cells were fixed in 4% PFA and stained with DAPI. The ratio of cells in the lower chamber versus total seeded cells was calculated.

### Wound healing assays

Cells were seeded in the appropriate number in a 6-well culture plates until 100% confluence was reached in 24 hours. Then, confluent cultures were scratched using a 200 μL pipette tip and then incubated at 37°C for 24 and 48 hours. At the indicated times, images of the wounds were acquired using the LEICA DMI6000 inverted microscope (Leica, Heidelberg, Germany) on a DC 350 FX camera (Leica). The wound areas were quantified using ImageJ software (NIH).

### Gelatin degradation (invasion) assay

Fluorescent-coated coverslips were prepared as described previously (Varone et al., 2021). Cells were plated on gelatin-coated coverslips in a 24-well plate and fixed after 24 and 48 hours with 4% PFA (v/v) for 15’ at room temperature. Then, filamentous actin and nuclei were stained using Alexa Fluor™ 488 Phalloidin (Invitrogen) and Hoechst 33342 (Invitrogen), respectively. Images were acquired using a confocal microscope (LSM 510; Zeiss) and degradation area were quantified using ImageJ software.

### Cell treatment

PDAC cells were seeded in a 6-well plate and after 24 hours were treated with 10 ng/ml of recombinant TGF-β1 (rTGF-β1, Peprotech) either alone or in the presence of 80 μM of Vactosertib (Vacto, MedChemExpress) every two days for a week. Cells were harvested by EDTA-trypsin, washed twice with PBS, and collected by centrifugation for RNA extraction.

### Cell growth and Chemoresistance assay

Proliferation rates were determined at different time points, as reported in the figures, using the CCK8 assay kit according to the manufacturer’s instruction (Dojindo; Kumamoto, Japan). For the chemoresistance assay, cells were treated with 100 μM of gemcitabine for 48 hours. Cell viability was determined using a the CCK-8 assay kit.

### Cell cycle assay

To synchronize the cell cultures, cells were seeded in 6 multiwell plate in growth medium with 10% FBS overnight. Then the cultures were rinsed by PBS and changed to serum free medium. After serum starvation for 24 hours, the cells were passaged and released into cell cycle by addition of serum. For FACS analysis, cell samples were harvested at indicated time points. Cells were trypsinized, washed in PBS, centrifuged, and pellets were fixed in 200 µl of 70% ethanol and stored at -20°C until use. Cells were centrifuged and pellets resuspended in 200 µl of PBS with 10 µg/mL of RNAse A. Cells were incubated for 1 hour at 37°C prior to resuspension in PI. Cell-cycle analysis was carried out by flow cytometry (CANTO II). Data were analyzed by FlowJo 9.2 software.

### Apoptosis assay

Attached and floating cells were collected, resuspended and stained with Annexin V (550474; BD Bioscience) after incubation with Annexin V binding buffer (556454, BD PharMingen). Cells were then incubated with PI. Samples were analyzed by flow cytometry using a FACS Canto II (BD), and data were analyzed using FlowJo 9.2 software.

### Tumor growth

All animal experiments have been approved by the local ministry (IACUC protocol #992/2017-PR) and were performed in the animal facility under pathogen-free conditions. Single-cell suspensions of 2.5×10^5^ L3.6pl, PANC-1 and #253 (sh empty or sh*LAMC2, EGFP*^*+*^ *or EGFP*^*-*^) cells were subcutaneously injected into 6-week-old nude CD1 male mice (Charles River Laboratories). Tumor take was monitored visually and by palpation bi-weekly. Tumor diameter and volume were calculated based on caliper measurements of tumor length and height using the formula: tumor volume = (length x width^2^)/2. Animals were considered to bear a tumor when the maximal tumor diameter was over 2 mm.

### Metastatic assay

Single-cell suspensions of 8×10^5^ #253 (EGFP^+^ or EGFP^-^) cells were injected into the tail vein of 6-week-old nude CD1 male mice (Charles River Laboratories) (n=5 per group). Vactosertib (40 mg/kg mice) was dissolved in DMSO and administered to mice two times *per* week for three weeks, by intraperitoneal injection. After 2 months from the injection, mice were sacrificed and lungs were resected and fixed in 4% PFA for hematoxylin and eosin staining.

### IHC in FFEPE

Immunostainings were carried out using 4-μm tissue sections according to standard procedures. Briefly, after antigen retrieval, samples were blocked with Peroxidase-Blocking Solution (Dako, S202386) for 10 min at room temperature, and primary antibodies were then incubated with samples overnight. Slides were washed with EnVision FLEX Wash Buffer (Dako, K800721), and the corresponding secondary antibody was incubated with the sample for 45 min at room temperature. Samples were developed using 3,3′-diaminobenzidine, counterstained with hematoxylin and mounted. Antibodies against LAMC2 (SIGMA, HPA024638-100U) and GFP (ABCA; ab290) were used at 1:100 dilution overnight at 4^°^C in the dark. The nuclei of cell were stained with Hematoxylin. Images were acquired using a digital image scanning (Nanozoomer 2.0HT, Hamamatsu) and cropped using NDP.view2. The area % stain represents the ratio of the summed absolute areas of staining versus the total tissue. The area % stain was analyzed by Fiji ImageJ version v3.2.28. The TMA (Biomax, PA484a) was stained following the above mentioned procedure. Images were acquired using a digital image scanning (Nanozoomer 2.0HT, Hamamatsu) and cropped using NDP.view2. The scoring algorithm takes the proportion of stained cells into consideration, as well as the intensity of the staining. The reactivity was scored in a semi-quantitative manner, which was categorized as low if less than 10% staining was observed in the epithelium; and medium or high based on the intensity if the percentage was between 10-25% and ≥25%, respectively.

### Histoscore (H-score)

The intensity of LAMC2 staining was reported based on the H-score method considering the intensity and the % of positive cells. We scored as: 1 for less than 5 %, 2 for 6-15 %, 3 for 15-39% and 4 for more than 40%.

### Donor plasmid construction

750 bp (LAMC2 construct) of 5’ homology arm (HA) and 3’ homology arm were amplified from PDAC gDNA and cloned in pDONOR vector. LF2A-EGFP-BGHpA insertion cassette was generated by gene synthesis (Thermo Fisher) and cloned in the 5’HA3’HA previously engineered pDONOR vector.

### sgRNA design

Small guide RNA was designed using the http://crispr.mit.edu web tool. To select for the most suitable sgRNAs, we applied the following criteria: i. localization of the sgRNA as near as possible to the desired site of insertion to maximize homologous recombination efficiency, ii. Cas9-mediated double strand break downstream of STOP codon to prevent NHEJ-induced indels in the ORF, iii. guide selected to anneal at the intersection between the 5’ homology arm and 3’ homology arm so that the donor plasmid is protected from Cas9 cut, iv. minimum off-target score according to http://crispr.mit.edu and maximum Doench activity score (Doench et al., 2014). sgRNAs used to modify PDAC cells was designed as follows: CCTCAGTTGAGAAATATTTATGG.

### px330-IRFP Cas9 plasmid construction

Px330 Cas9 plasmid from Feng Zhang’s laboratory was obtained from addgene (ref. 42230) and was modified by the introduction of a SV40promoter-IRFP expression cassette downstream of Cas9 by FseI – EcoRI. In addition the BbsI site of IRFP was silenced by site-directed mutagenesis. SgRNAs were cloned in px330-IRFP as described in www.genome-engineering.org/crispr/wpcontent/uploads/2014/05/CRISPR-Reagent-Description-Rev20140509.pdf

### Nucleofection

For PDAC cells nucleofection, one million of single-cell trypsinized PDAC cells were nucleofected with 3 µg of donor plasmid and 1 µg of px330-IRFP Cas9 corresponding plasmids using Lonza nucleofector kit V (VVCA-1003) and program A-32 in an Amaxa-II nucleofector following manufacturer protocol.

### FACS strategy and generation of single cell-derived PDAC cells

Nucleofected cells were cultured in DMEM/RPMI complete medium for 48 hours. Then, we isolated cells that were IRFP positive by FACS and we cultured them in complete medium for 20 days we sorted the cells for EGFP to confirm the donor integration. We then selected the cell population positive and negative for the expression of EGFP (LAMC2-LF2A-EGFP) and we named them EGFP^+^ and EGFP^-^, respectively. Cells were seeded in a 96-well format to derive single-cell clones.

### Specific genotyping PCRs

Single-cell derived clones were lysed in buffer consisting of 10 mM Tris, 1 mM EDTA, 1 % Tween 20 and 0.4 mg/ml proteinase K for 1 h at 55 °C. The lysate was directly used in the specific integration PCR. For the 5’ specific integration PCR a forward primer upstream of the 5’ homology arm and a reverse primer at the beginning of the inserted cassette were used. Similarly, for the 3’ specific integration PCR a forward primer at the end of the inserted cassette and a reverse primer downstream of the 3’ homology arm were used. The PCR conditions were as follows: DNA Polymerase (BioTools #10012-4103) 95 °C 2 min x38 (95 °C 30 s – 55 °C 30 s – 72 °C 1:30 min) 72 °C 5 min - hold 16 °C. Used primer sequences are shown in **Table 2**.

**Table 2:**
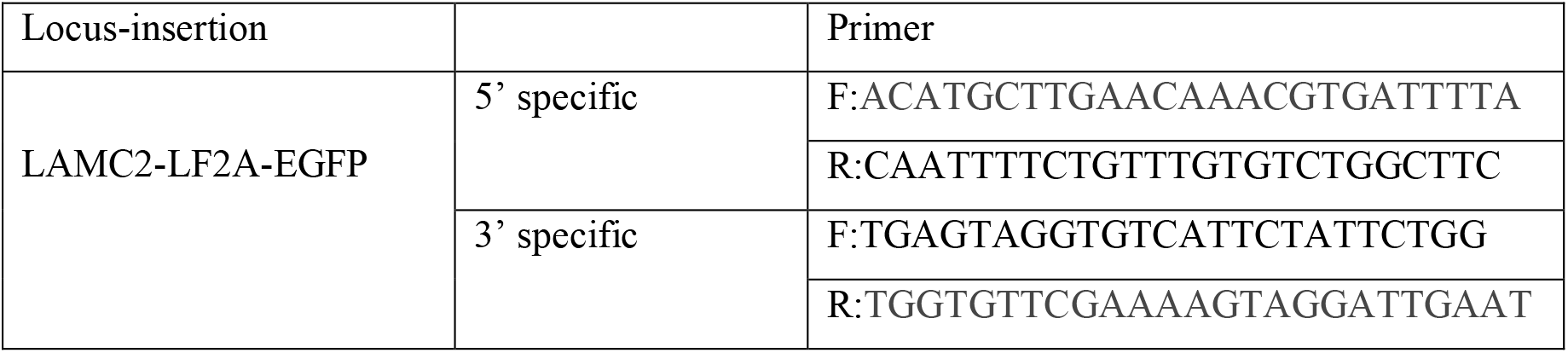
Primers used for the specific integration PCR.

### Bioinformatics analysis

Normalized expression data was downloaded from NCBI GEO (GSE21501, GSE28735, GSE62452 and GSE71729) with the R package GEOquery. GSE621501 consists of 118 PDAC samples and 13 control samples. GSE28735 consists of 45 tumor samples and matching normal samples. GSE62452 consists of 69 PDAC tumors and 61 normal pancreatic tissue samples from pancreatic cancer patients. GSE71729 consists of 145 primary and 61 metastatic PDAC tumors, 17 cell lines, 46 pancreas and 88 distant site adjacent normal samples A Principal Component Analysis was performed on the entire dataset, which showed a clear separation between tumor and non-tumor samples. Survival was analyzed using the http://gepia2.cancer-pku.cn/#survival. A Median Group cutoff (50% high vs 50% Low) was used. The analysis was performed considering only the classical PAAD subtype (84 patients). The Pearson’s Correlation analysis was performed using the data reported in the http://www.analytics.pancreasexpression.org/index.php?s=icgc. The TCGA dataset is composed by 84 patients with PDAC, the USA cohort is composed by 185 patients with PDAC and the Canadian (CA) cohort is composed by 317 patients with Pancreatic Cancer.

### Gene expression data sets and GSEA analyses

The gene expression data sets used in this study are publicly available. The dataset from Jandaghi *et al*. (Jandaghi et al., 2016) was downloaded from ArrayExpress (E-MTAB-1791); the data set from Janky *et al*. (Janky et al., 2016) was downloaded from GEO (GSE62165); the dataset from Bailey et al. was included in a supplementary figure of their published work (Australian Pancreatic Cancer Genome Initiative et al., 2016); the META data set, containing data sets GSE15471, GSE16515, GSE22780, and GSE32688, was generated as described in (Martinelli et al., 2017) and the TCGA dataset was downloaded from The Cancer Genome Atlas (TCGA; http://xena.ucsc.edu). The samples included in the top and bottom quartile of expression of LAMC2 were compared in GSEA, using the Hallmark gene-set database. The GSEA module of the Genepattern suite from the Broad Institute was used, with 1000 permutations and FDR < 25% was considered statistically significant.

### RNA-sequencing analysis

Total RNAs were extracted with Eurogold TRIFAST kit (Euroclone) according to the manufacturer’s instructions. TruSeq stranded mRNA libraries were run on the Illumina NextSeq 500. We generated 40 million reads per lane and paired-end reads from sequencing were trimmed from adaptors and quality checked by the sequencing service. Reads were aligned against the human reference sequence GRCh38 (hg38) from Genome Reference Consortium using BWA (Li 2009). Read counts per gene were determined using the feature Counts method from the Subread R package (Liao 2014) based on the gene information in the gtf available here ftp://ftp.ensembl.org/pub/release-94/gtf/homo_sapiens/Homo_sapiens.GRCh38.94.gtf.gz. Gene signatures (Hallmark genesets) were downloaded from GSEA—Molecular Signature Database for Gene set enrichment analysis. RNA-seq data for #253-LAMC2-EGFP high vs. negative cells is available at https://www.ebi.ac.uk/arrayexpress/ under accession number E-MTAB-11597.

### Statistical Analyses

Results for continuous variables are presented as means ± standard deviation (SD) unless stated otherwise of at least three independent experiments. Treatment groups were compared with the independent samples t test. Pair-wise multiple comparisons were performed with the one-way ANOVA (two-sided) with Bonferroni adjustment. The disease-free interval of patients was calculated using the Kaplan– Meier method, and differences among subgroups were assessed by the log-rank test. Experiments were performed a minimum of three independent times and always performed in independent triplicate samples. qPCR were repeated a minimum of five independent times in triplicate. p < 0.05 was considered statistically significant. All analyses were performed using GraphPAD Prism7. Correlation analysis were performed applying the Pearson’s correlation coefficient.

## ACKNOWLEDGEMENTS

We are grateful to members of the Integrated Microscopy and FACS Facilities of IGB-ABT and to the EuroBioImaging Infrastructure at Institute for Experimental Endocrinology and Oncology (CNR), Naples. The human immortalized HPNE were kindly provided by Dr. Francisco X Real. This work was supported by the Marie Curie IF (H2020-MSCA-IF-2015, #703753), My First AIRC Grant (MFAG-2017, #20206), POR Campania FESR 2014/2020 (Project SATIN) to E.L., AIRC IG grant 2018 n.21420 to A.D.L. and FIMP to D.D.C.

## AUTHOR CONTRIBUTIONS

EL designed the experiments. DDC, ADD and TTI performed and analyzed the experiments. BS, SB and VC analyzed the RNA-seq data. CCD designed the CRISPR/Cas9 cloning strategy. MS performed the IHC on the TMA slide. MC and AC performed the *in vivo* experiments. CH and BS provided patient specimens. EL assembled results and wrote the paper. All authors participated in critical revision of the manuscript for important intellectual content.

