## Supplemental Data for "LAMC2 marks a tumor-initiating cell population with an aggressive signature in pancreatic cancer"

**Supplementary Figure 1 – Increased LAMC2 expression is associated with unfavorable outcome in PDAC**

(**a**) Representative images of IHC staining for LAMC2 (brown) in tissue section from pancreatic islet. (**b**) *H*-score for LAMC2 expression. (**c**) Number of PDAC tumors (Human Protein Atlas database) classified based on LAMC2 immunohistochemistry for staining, intensity and quantity. (**d**) Patients distribution according to gender, age and TNM. (**e**) *H-score* and grade distribution according to gender and age. (**f**) Kaplan-Meier curves showing overall survival of PDAC patients, stratified according to the median value of LAMC2 expression for gender, alcohol consumption and smoking.

**Supplementary Figure 2 – LAMC2 expression correlates with CSC content and function**

(**a**) qPCR analysis for *LAMC2* gene in adherent cells versus spheres. Data are normalized to *GAPDH* and are presented as fold change in gene expression relative to the adherent counterpart (indicated as Adh). (**b**) Western blot analysis of LAMC2 in adherent cells versus spheres. Parallel β-ACTIN immunoblotting was performed. (**c**) qPCR analysis for *CD44, CD133* and *L1CAM* genes in in adherent cells versus spheres. Data are normalized to *GAPDH* and are presented as fold change in gene expression relative to adherent cells. (**d)** qPCR analysis for *LAMC2* in PDAC cells grown in different culture conditions. (**e**) qPCR analysis for *LAMC2, ABCG2, CD133, L1CAM* and *SOX2* in Adh *vs* Spheres *vs* Differentiated cells. Data are normalized to *GAPDH* and are presented as fold change in gene expression relative to adherent cells. (**f**) qPCR analysis for *CD44* and *LAMC2* genes in $CD44^+$ sorted cells. Data are normalized to *GAPDH* and are presented as fold change in gene expression relative to $CD44^-$ cells. (**g**) qPCR analysis for *CD133* and *LAMC2* genes in $CD133^+$ sorted cells. Data are normalized to *GAPDH* and are presented as fold change in gene expression relative to $CD133^-$ cells. (**h**) qPCR analysis for *LAMC2* in PDAC cells treated with 100 µM of gemcitabine. Data are normalized to *GAPDH* and are presented as fold change in gene expression relative to control cells (untreated). $^*p<0.05$, $^{**}p<0.005$, $^{***}p<0.0005$. n≥ 3.

**Supplementary Figure 3 – Knockdown of LAMC2 does not affect cell growth**

(**a**) Western blot analysis of LAMC2 in sh empty and *LAMC2* knockdown cells. Parallel β-ACTIN immunoblotting was performed. (**b**) qPCR analysis for *LAMC2* and CD44 genes in sh empty and *LAMC2* knockdown cells. Data are normalized to *GAPDH* and are presented as fold change in gene expression relative to sh empty. (**c**) Representative images of sh empty and *LAMC2* knockdown cells grew in monolayer. (**d**) Cell viability of sh empty and *LAMC2* knockdown cells. Cell viability was evaluated using cell-counting-kit 8, and absorbance was measured at 450 nm. (**e**) Cell cycle analysis of sh empty and *LAMC2* knockdown cells. (PI incorporation). $^*p<0.05$, $^{**}p<0.005$, $^{***}p<0.0005$. n≥ 3.

**Supplementary Figure 4 – Loss of LAMC2 reduces stemness**

(**a**) Flow cytometry for apoptotic cells as determined by AnnexinV/PI staining in control and *L1* knockdown cells. (**b**) Flow cytometry quantification of CD44 in sh empty and *LAMC2* knockdown cells. (**c**) qPCR analysis for *CD133* gene in sh empty and *LAMC2* knockdown cells. Data are normalized to *GAPDH* and are presented as fold change in gene expression relative to sh empty. (**d**) Flow cytometry quantification of CD133 in sh empty and *LAMC2* knockdown cells. (**e**) Representative images of sh empty and *LAMC2* knockdown cells grew in spheres. (**f**) Sphere formation capacity of sh empty and *LAMC2* knockdown cells. P1= 1$^{st}$ generation; P2= 2$^{nd}$ generation. (**g**) Quantification of sphere size of sh empty and *LAMC2* knockdown cells. (**h**) Organoid formation capacity of sh empty and *LAMC2* knockdown cells. (**i**) Migration assay of sh empty and *LAMC2* knockdown cells. The nuclei were stained in blue (DAPI). (**j**) Migratory potential of sh empty and *LAMC2* knockdown cells. (**k**) Wound healing assay of sh empty and *LAMC2* knockdown cells. *$p<0.05$, **$p<0.005$, ***$p<0.0005$. $n \geq 3$.

**Supplementary Figure 5 – Knockdown of LAMC2 affects tumorigenicity**

(**a**) Representative images of gelatin degradation assay of sh empty and *LAMC2* knockdown cells. Nuclei were stained with Hoechst 33342 (blue), green represents actin (Alexa Fluor™ 488 Phalloidin) and red illustrates gelatin (Rodhamine). (**b**) Degradation potential of sh empty and *LAMC2* knockdown cells. (**c**) qPCR analysis for EMT genes in sh empty and *LAMC2* knockdown cells. Data are normalized to *GAPDH* and are presented as fold change in gene expression relative to sh empty. (**d**) Western blot analysis of VIM in sh empty and *LAMC2* knockdown cells. Parallel β-ACTIN immunoblotting was performed. (**e**) Western blot analysis of MMP2 in sh empty and *LAMC2* knockdown cells. Parallel β-ACTIN immunoblotting was performed. (**f**) Number of tumors generated by the subcutaneous injection of sh empty and *LAMC2* knockdown cells. (**g**) qPCR analysis for *LAMC2*, EMT, *MMP2* and *MMP10* genes in sh empty and *LAMC2* knockdown cells isolated from tumors. Data are normalized to *GAPDH* and are presented as fold change in gene expression relative to sh empty. *$p<0.05$, **$p<0.005$, ***$p<0.0005$. $n \geq 3$.

**Supplementary Figure 6 – Generation of *LAMC2*-EGFP knock-in human PDAC cells**

(**a**) Flow cytometry for IRFP in PDAC cells 48 hours post-nucleofection. All cytometry gates were established based on isotype controls. (**b**) Flow cytometry for EGFP in PDAC cells 20 days post-nucleofection. All cytometry gates were established based on isotype controls. (**c**) Specific integration PCR gDNA analysis. The position of primers is indicated by arrows. (**d**) FACS profiles showing the expression of EGFP in the EGFP$^-$ and EGFP$^+$ cells. (**e**) qPCR analysis for *LAMC2* gene in EGFP$^+$ and EGFP$^-$ cells. Data are normalized to *GAPDH* and are presented as fold change in gene expression relative to the EGFP$^-$ counterpart. (**f**) Representative images of EGFP$^+$ and EGFP$^-$ cells grew in monolayer. (**g**) Cell cycle analysis of EGFP$^+$ and EGFP$^-$ cells (PI incorporation). (**h**) Sphere formation capacity of EGFP$^+$ and EGFP$^-$ cells. *$p<0.05$, **$p<0.005$, ***$p<0.0005$. $n \geq 3$.

**Supplementary Figure 7 – Characterization of human *LAMC2*-EGFP PDAC cells *in vitro* and *in vivo***

(**a**) Representative images of EGFP$^+$ and EGFP$^-$ cells grew in Matrigel. (**b**) Organoid formation capacity of EGFP$^+$ and EGFP$^-$ cells. (**c**) Quantification of organoids size of EGFP$^+$ and EGFP$^-$ cells. (**d**) qPCR analysis for

*LAMC2*, EMT, *MMP2* and *MMP10* genes in $EGFP^+$ and $EGFP^-$ cells. Data are normalized to *GAPDH* and are presented as fold change in gene expression relative to the $EGFP^-$ counterpart. (**e**) Tumor volume of $EGFP^+$ and $EGFP^-$ cells subcutaneously injected into athymic mice. n ≥ 10. (**f**) Representative histologic sections of xenografts derived from $EGFP^+$ and $EGFP^-$. The tumor sections were stained for H&E and EGFP. (**g**) Representative flow cytometry for EGFP in subcutaneous tumors derived from $EGFP^+$ and $EGFP^-$ cells. (**h**) qPCR analysis for *LAMC2*, EMT, *MMP2* and *MMP10* genes in $EGFP^+$ and $EGFP^-$ cells isolated from tumors. Data are normalized to *GAPDH* and are presented as fold change in gene expression relative to the $EGFP^-$ counterpart. (**i**) qPCR analysis for *CDH1* gene in $EGFP^+$ and $EGFP^-$ cells isolated from tumors. Data are normalized to *GAPDH* and are presented as fold change in gene expression relative to the $EGFP^-$ counterpart. *p<0.05, **p<0.005, ***p<0.0005. n≥ 3.

**Supplementary Figure 8 – Global gene expression profiles of LAMC2-$EGFP^+$ and $EGFP^-$-derived tumors**

Heat map of differentially expressed genes in $EGFP^+$ and $EGFP^-$ cells isolated from tumors.

**Supplementary Figure 9 – Transforming growth factor beta (TGF-β) signaling inhibitor blocks LAMC2-induced metastasis**

(**a**) Enrichment plots for pancreatic cancer, focal adhesion, hypoxia, MAPK signaling, glycolysis and gluconeogenesis pathways in $EGFP^+$ versus $EGFP^-$ cells isolated by FACS from subcutaneous tumors. (**b**) Representative immunofluorescence images for LAMC2 (red) and nuclei (blue, DAPI) of PDAC cells treated with 10 ng/ml of recombinant TGF-β (rTGF-β1 and rTGF-β2). (**c**) Quantification of lung metastasis area on H&E-stained sections. *p<0.05, n≥ 5.

**Supplementary Figure 1_Increased LAMC2 expression is associated with unfavorable outcome in PDAC**

a

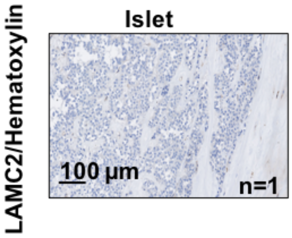


b

| | H-score | | | |
|---|---|---|---|---|
| | 1 (<5%) | 2 (6-15%) | 3 (15-39%) | 4 (>40%) |
| Islet | 100% | 0% | 0% | 0% |

c

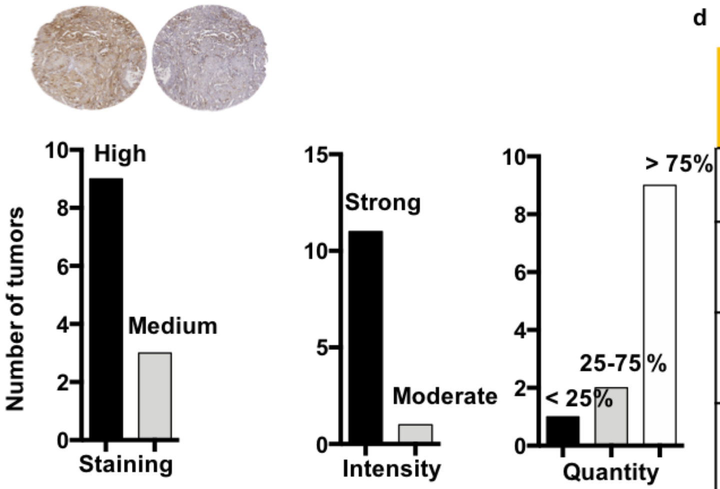


d

| Variable tested | | Number of patients |
|---|---|---|
| Group | PDAC | 20 |
| Gender | Male | 11 |
| | Female | 9 |
| Age | <60 | 11 |
| | >60 | 9 |
| Depth of tumor invasion (T factor) | T2 | 5 |
| | T3 | 14 |
| | T4 | 1 |
| Nodal involvement (N factor) | N0 | 16 |
| | N1 | 4 |
| Distant metastases (M factor) | No distant metastases | 20 |

e

| Variable tested | | Number of patients | | | |
|---|---|---|---|---|---|
| | | Female | Male | Age <60 | Age >60 |
| H-score | <30% | 7 | 5 | 6 | 7 |
| | >30% | 2 | 6 | 5 | 2 |
| Grade | G1 | 4 | 1 | 3 | 2 |
| | G2 | 3 | 8 | 6 | 5 |
| | G3 | 2 | 2 | 2 | 2 |

f

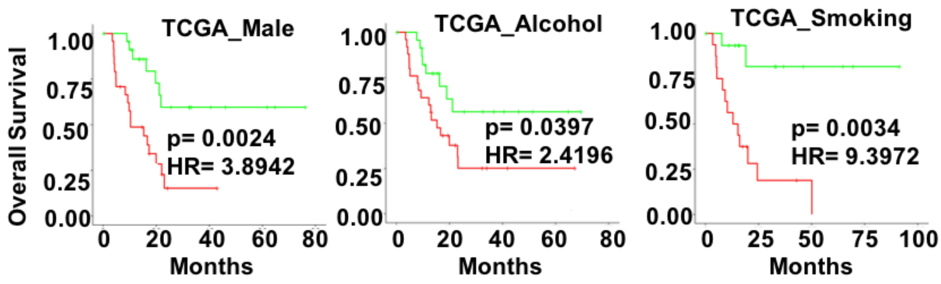

Supplementary Figure 2_LAMC2 expression correlates with CSC content and function

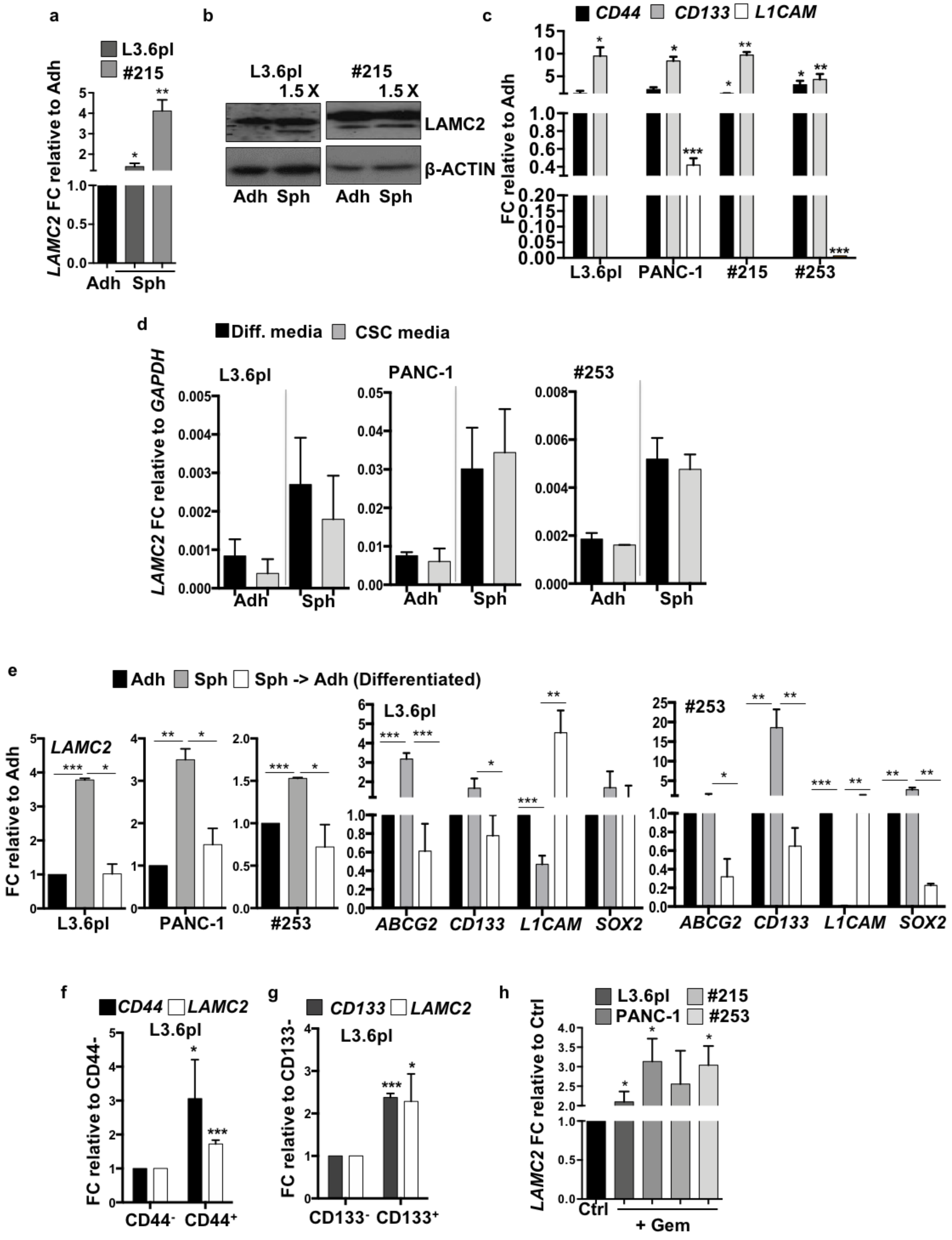

**Supplementary Figure 3_Knockdown of LAMC2 does not affects cell growth**

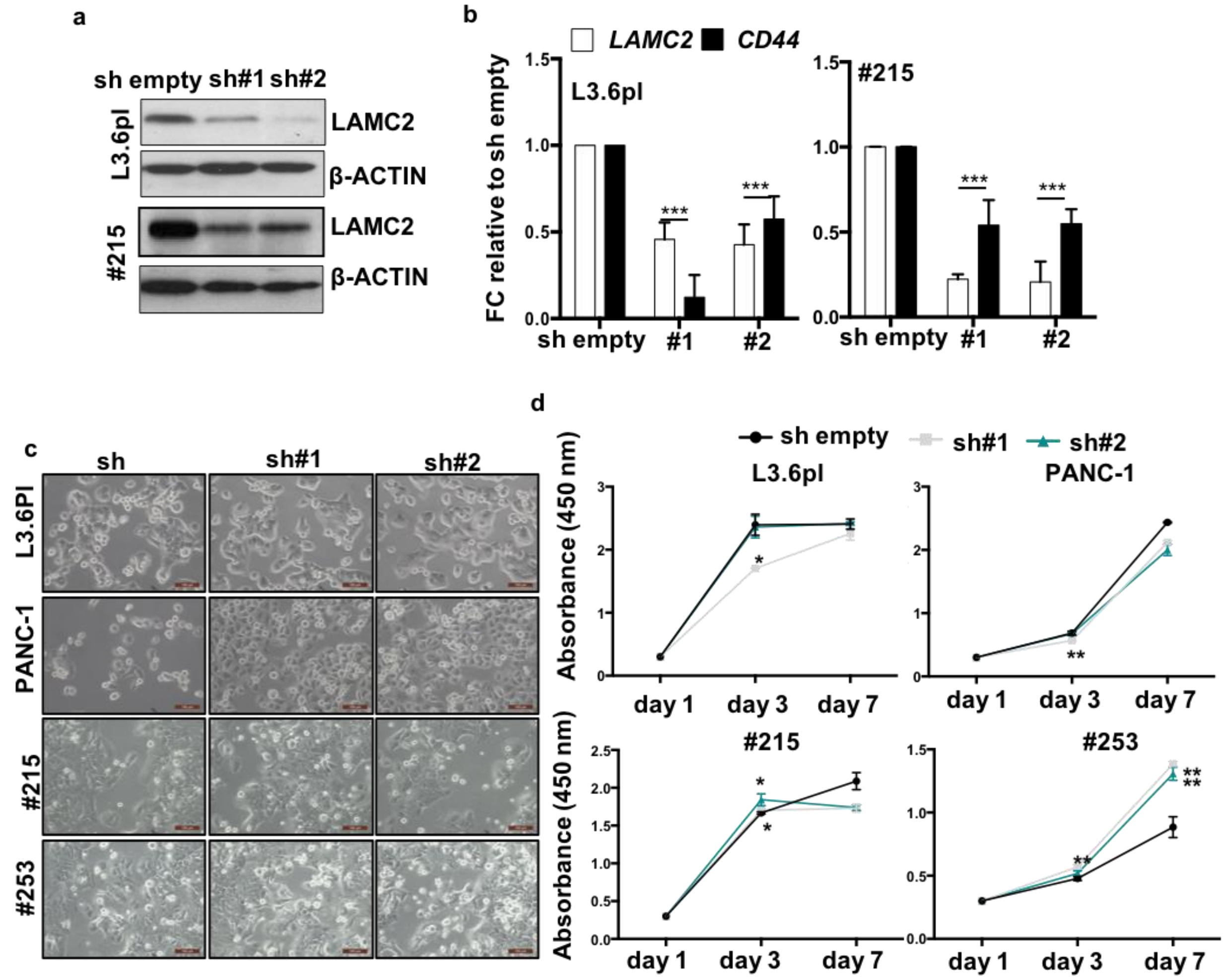


e

| Cell Cycle 48h | | pre G0 (%) | G0/G1 (%) | S (%) | G2/M (%) |
|---|---|---|---|---|---|
| L3.6pl | sh empty | 2.4 | 75.9 | 9 | 12.7 |
| | sh#1 | 2.7 | 71.8 | 11.1 | 14.4 |
| | sh#2 | 12.9 | 62.6 | 11.6 | 12.7 |
| PANC-1 | sh empty | 0.55 (± 0.07) | 40.75 (± 5.87) | 17.1 (± 2.12) | 41.3 (± 3.54) |
| | sh#1 | 0.85 (± 0.92) | 48.2 (± 3.25) | 21.2 (± 2.40) | 29.6 (± 1.70) |
| | sh#2 | 0.55 (± 0.49) | 49.2 (± 3.54) | 21.05 (± 2.33) | 29.15 (± 1.63) * |
| #215 | sh empty | 0.5 | 44.9 | 15.6 | 38.8 |
| | sh#1 | 1.5 | 50.5 | 19.5 | 28.4 |
| | sh#2 | 0.9 | 51.7 | 19.4 | 28 |
| #253 | sh empty | 4.55 (± 0.92) | 57.6 (± 2.12) | 19.15 (± 1.91) | 18.6 (± 1.13) |
| | sh#1 | 6.4 (± 2.55) | 58.35 (± 1.48) | 15.85 (± 1.20) | 19.25 (± 5.30) |
| | sh#2 | 6.75 (± 4.45) | 57.4 (± 2.55) | 15.35 (± 3.18) | 20.4 (± 5.09) |

**Supplementary Figure 4_Loss of LAMC2 reduces stemness**

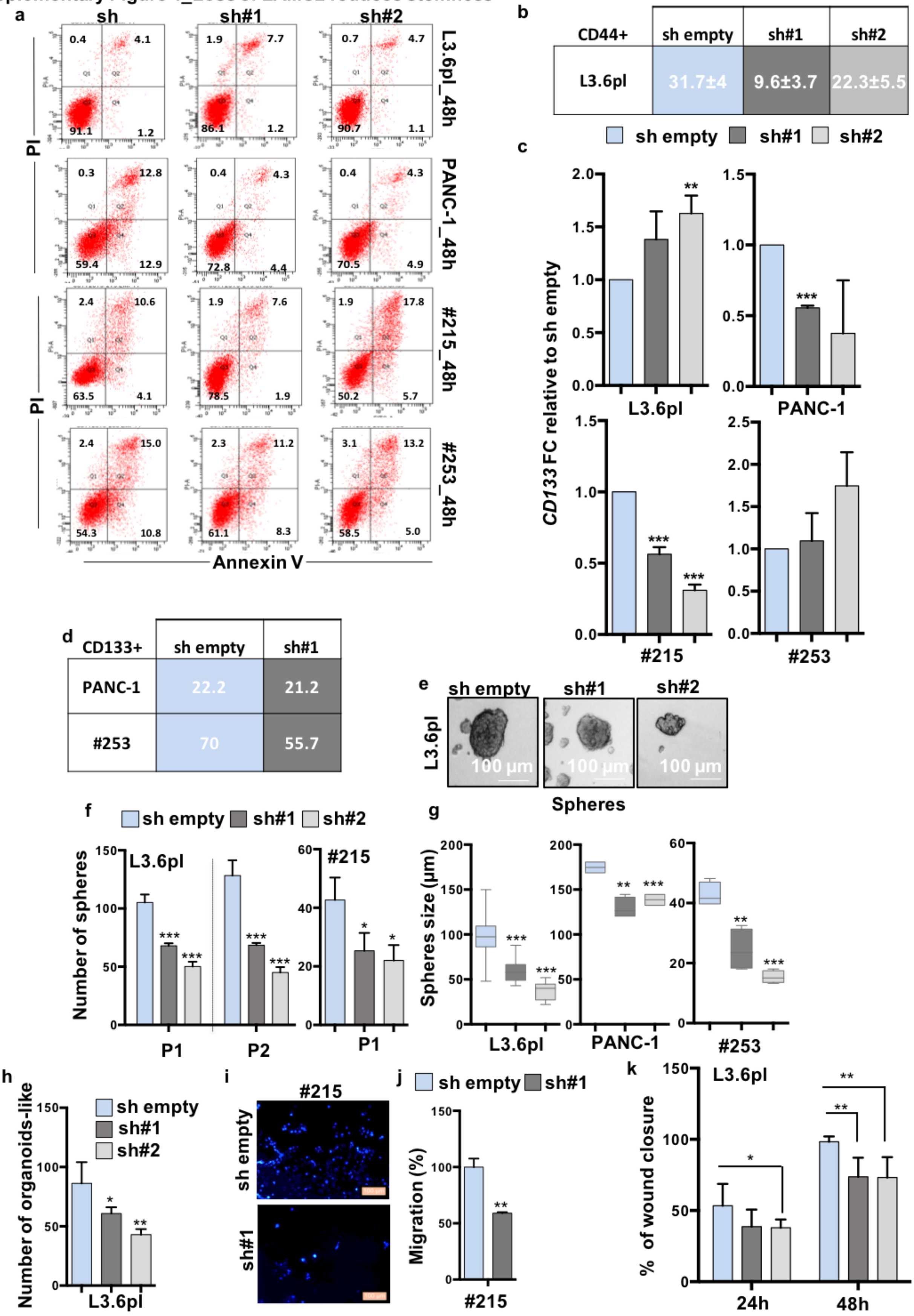

Supplementary Figure 5_Knockdown of LAMC2 affects tumorigenicity

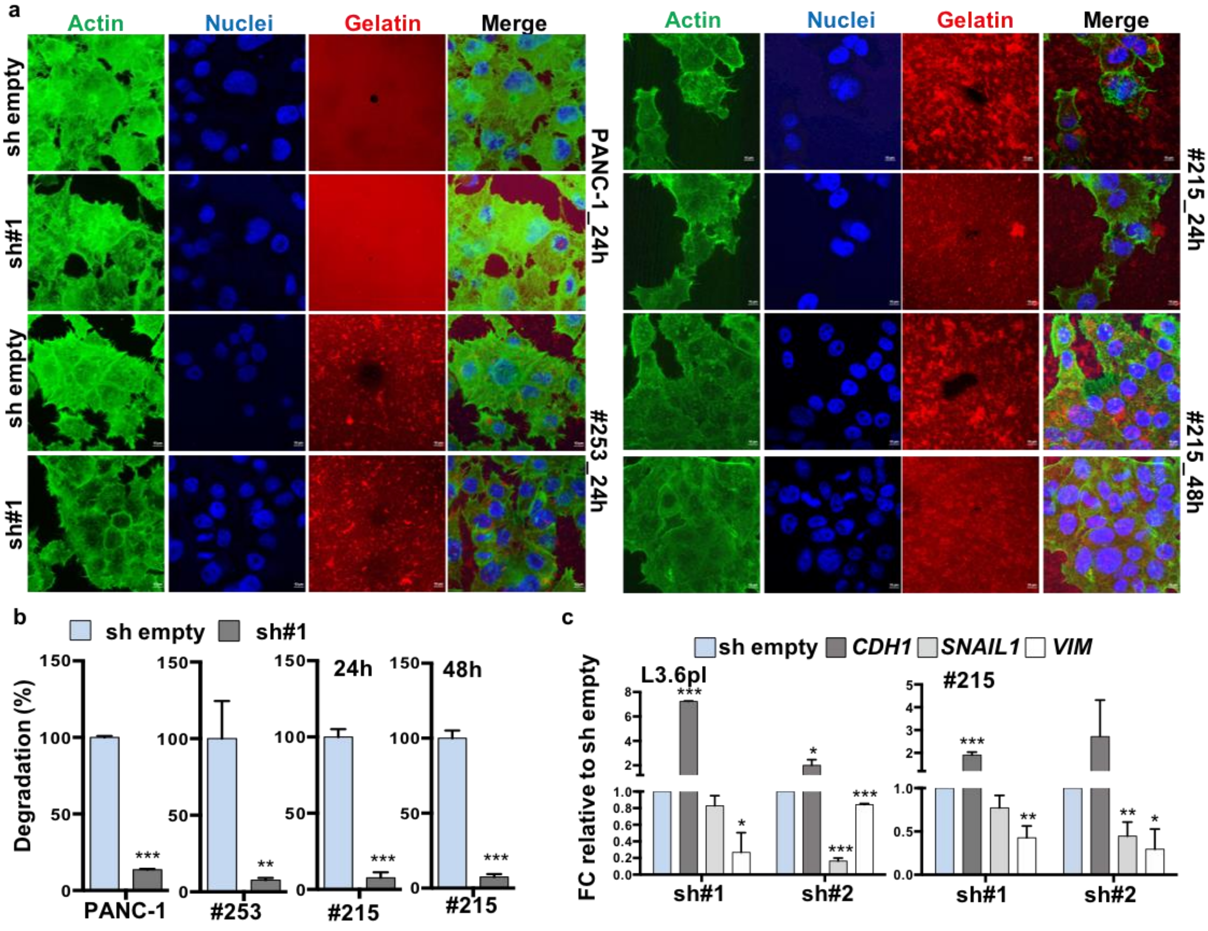


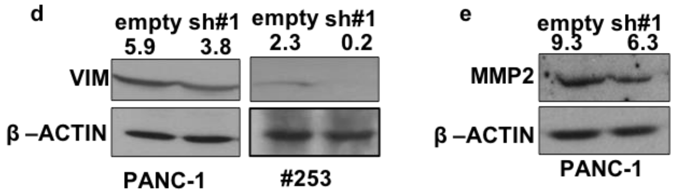


f

| Cells | Number of tumors |
|---|---|
| L3.6pl sh empty | 6/6 |
| L3.6pl sh#1 | 3/6 |
| PANC-1 sh empty | 7/8 |
| PANC-1 sh#1 | 1/8 |
| 215 empty | 2/6 |
| 215 sh#1 | 1/6 |
| 253 empty | 3/6 |
| 253 sh#1 | 3/6 |

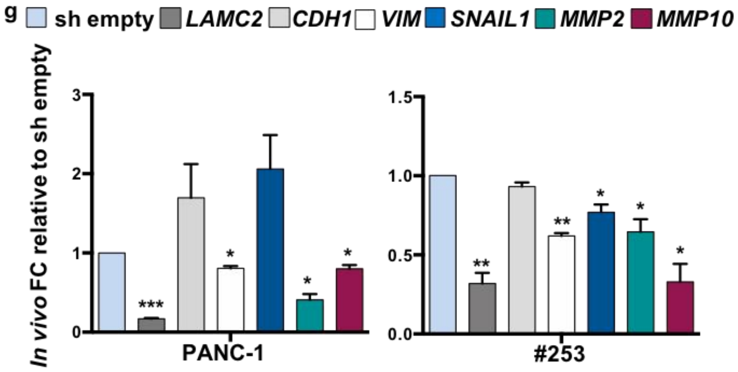

**Supplementary Figure 6_ Generation of *LAMC2*-EGFP knock-in human PDAC cells**

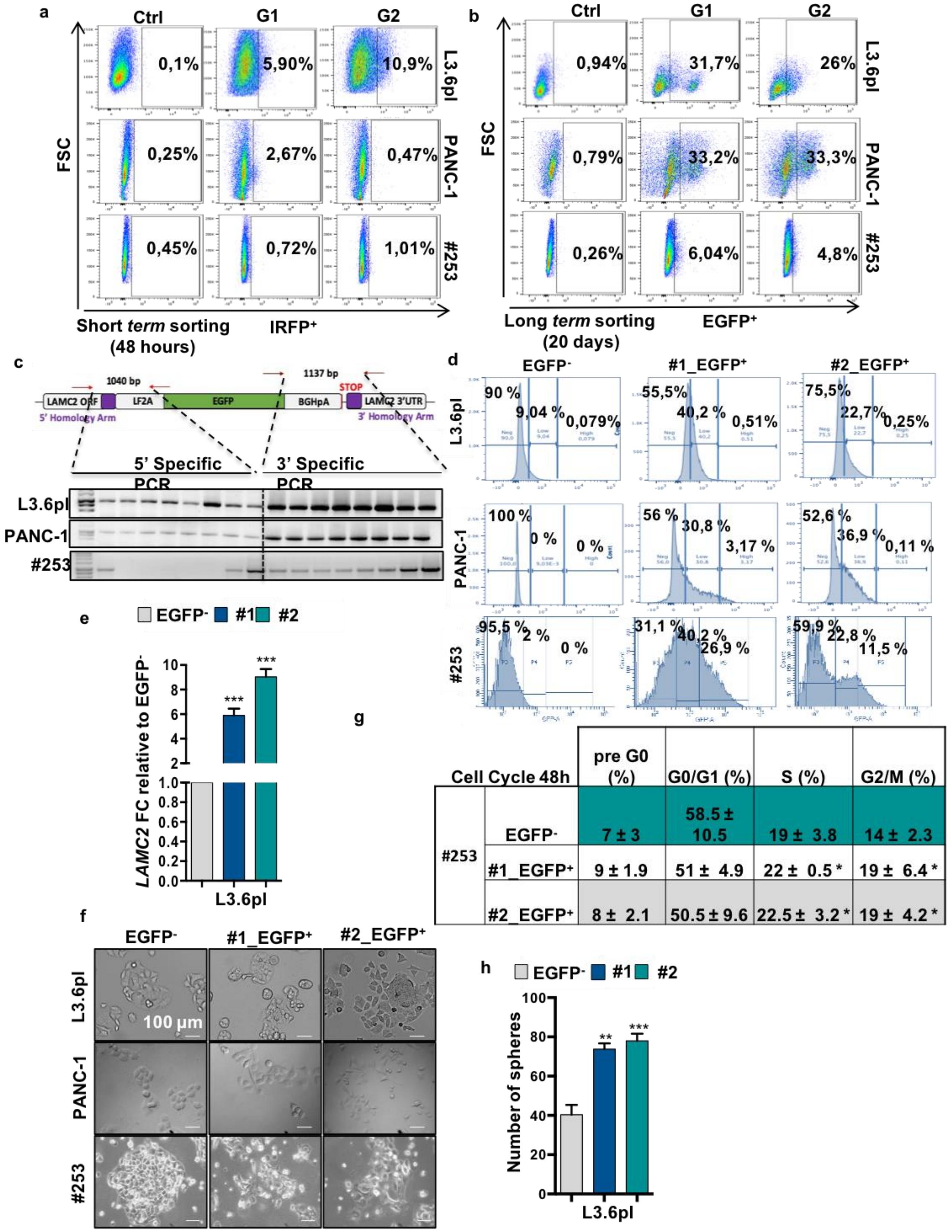


| Cell Cycle 48h | | pre G0 (%) | G0/G1 (%) | S (%) | G2/M (%) |
|---|---|---|---|---|---|
| #253 | EGFP- | 7 ± 3 | 58.5 ± 10.5 | 19 ± 3.8 | 14 ± 2.3 |
| | #1_EGFP+ | 9 ± 1.9 | 51 ± 4.9 | 22 ± 0.5 * | 19 ± 6.4 * |
| | #2_EGFP+ | 8 ± 2.1 | 50.5 ± 9.6 | 22.5 ± 3.2 * | 19 ± 4.2 * |

Supplementary Figure 7_Characterization of human *LAMC2*-EGFP PDAC cells *in vitro* and *in vivo*

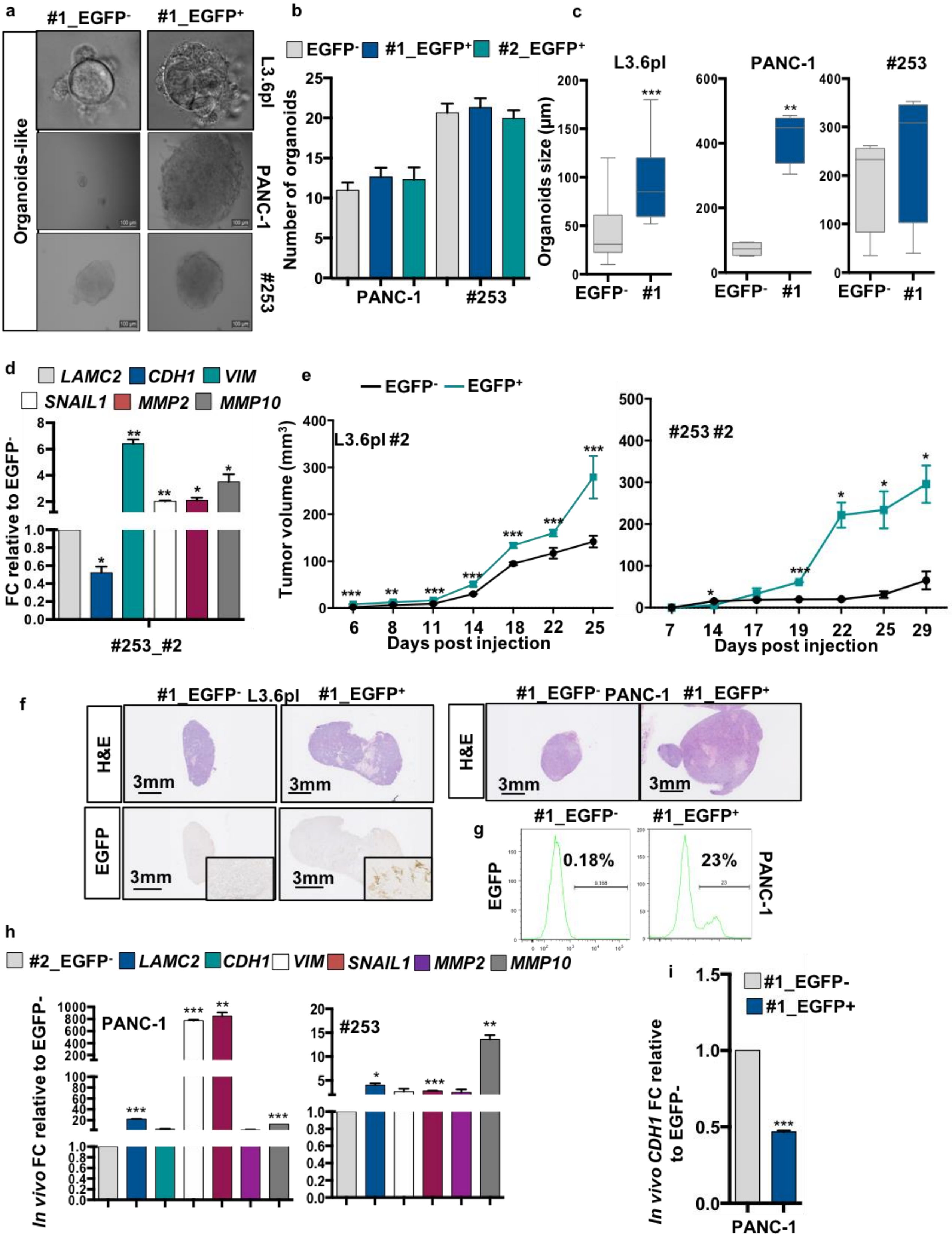

**Supplementary Figure 8_Global gene expression profiles of LAMC2-EGFP$^{+}$ and EGFP$^{-}$-derived tumors**

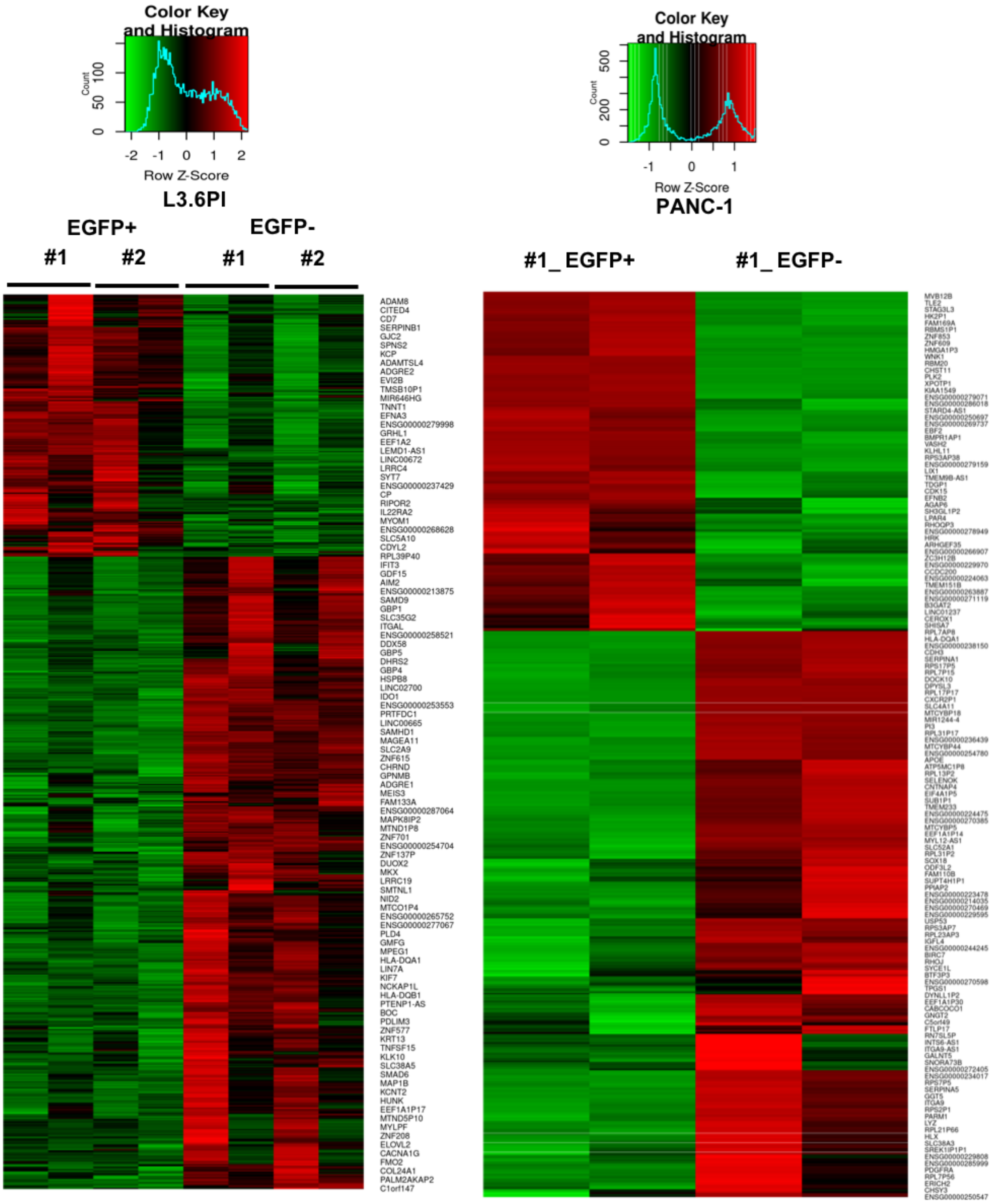

**Supplementary Figure 9_Transforming growth factor beta (TGF-β) signaling inhibitor blocks LAMC2-induced metastasis**

a

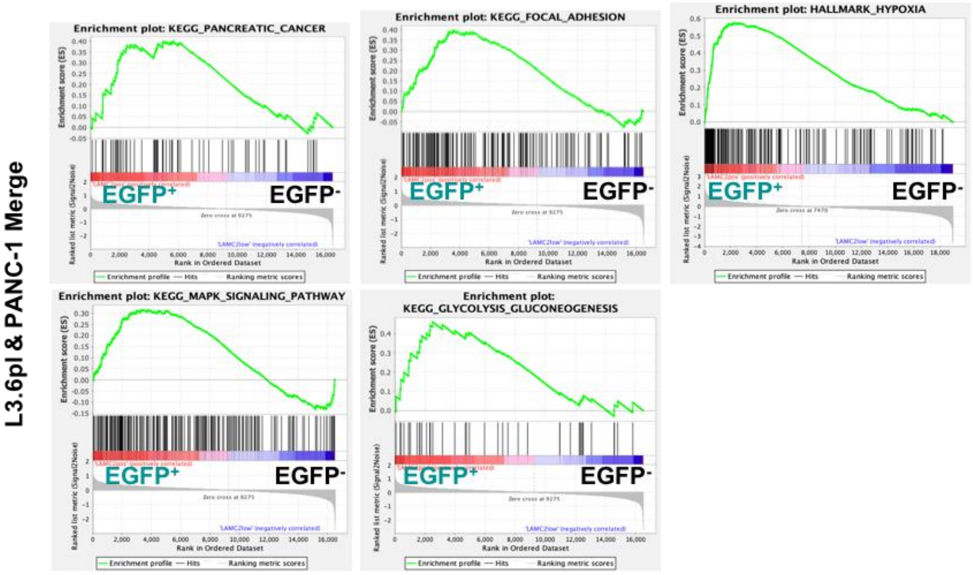


b

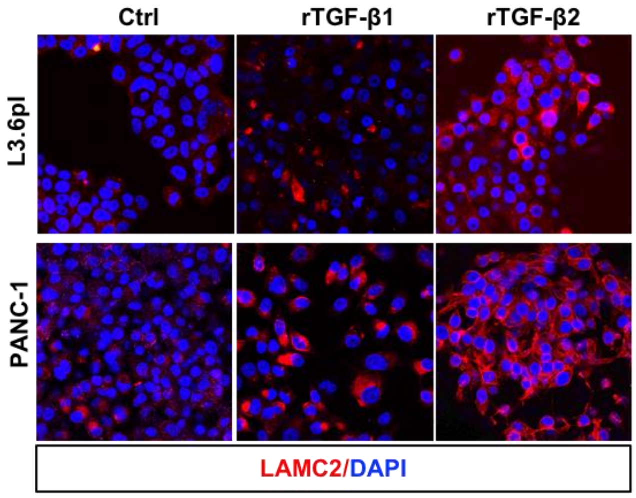


c

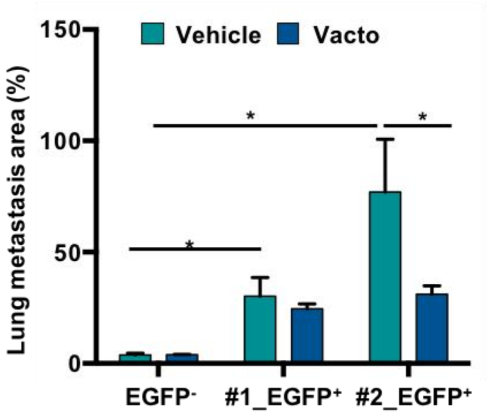
